# Non-canonical Activation of Human Group 2 Innate Lymphoid Cells by TLR4 Signaling

**DOI:** 10.1101/2020.10.29.361345

**Authors:** Li She, Hamad H. Alanazi, Jingwei Wang, Daniel P. Chupp, Yijiang Xu, Hong Zan, Zhenming Xu, Yilun Sun, Na Xiong, Nu Zhang, Xin Zhang, Yong Liu, Xiao-Dong Li

**Affiliations:** Department of Microbiology, Immunology and Molecular Genetics Long, School of Medicine, University of Texas Health San Antonio, San Antonio, TX 78229-3900; Department of Otolaryngology-Head and Neck Surgery, Central South University Changsha, Hunan 410008, China; Clinical Research Center for Pharyngolaryngeal Diseases and Voice Disorders, Central South University, Changsha, Hunan 410008, China; Otolaryngology Major Disease Research Key Laboratory of Hunan Province, Central South University, Changsha, Hunan 410008, China; National Clinical Research Center for Geriatric Disorders, Xiangya Hospital, Central South University, Changsha, Hunan 410008, China; Institute of Pediatrics, The Second Xiangya Hospital, Central South University, Changsha, Hunan 410008, China

**Keywords:** group 2 innate lymphoid cells (ILC2), toll-like receptor 4 (TLR4), lipopolysaccharides (LPS), interleukin 33 (IL-33), Interleukin 1 receptor-like 1 (IL1RL1, also called IL-33R or ST2), interleukin 4 (IL-4), interleukin 5 (IL-5), interleukin 13 (IL-13)

## Abstract

Group 2 innate lymphoid cells (ILC2) are emerging as a critical player in type 2 immunity at barrier sites in response to microbial infections and allergen exposures. Although their classical activators are known to be host epithelial-derived alarmin cytokines IL-33, IL-25 or TSLP, it remains elusive whether ILC2 cells can be activated by directly sensing microbial ligands via pattern-recognition receptors such as toll-like receptors (TLRs). Here we report that toll-like receptor 4 (TLR4) is a potent activating receptor of human ILC2. We found that among many microbial ligands examined, lipopolysaccharides (LPS) from multiple species of Gram-negative bacteria, was found to potently stimulate human, but not murine ILC2, to proliferate and produce massive amounts of type 2 effector cytokines IL-4, IL-5, and IL-13. LPS-activated ILC2 also had greatly enhanced the CD40 ligand (CD154) expression and were able to promote the proliferation and antibody production of human B cells in culture. In a humanized mouse model, LPS activated the adoptively transferred human ILC2 in mouse lungs. Both NF-kB and JAK pathways, but not the IL-33-ST2 pathway, were required for LPS to activate human ILC2. RNA-seq data further revealed that LPS induced a large set of genes overlapped significantly with those induced by IL-33. Collectively, these findings support a non-classical mode of activating human ILC2 cells via the LPS-TLR4 signaling axis. Thus, targeting TLR4 signaling pathway might be developed as a new approach by modulating ILC2 activation in treating various type 2 immunity-associated diseases.

## Introduction

Allergic disorders including asthma, allergic rhinitis and atopic dermatitis (AAA) are common diseases affecting more than 300 million people worldwide (Jackson et al., 2013; Kay, 2001a; Kay, 2001b). Although AAA diseases are traditionally characterized by an overzealous Th2-mediated inflammatory response, it has become increasingly appreciated in recent years that group 2 innate lymphoid cells (ILC2) play an important role in the initiation and orchestration of type 2 immunopathologies (Vivier et al., 2018). ILC2 cells are the innate counterparts of Th2 lymphocytes, but lack rearranged antigen receptors (Artis and Spits, 2015; Barlow and McKenzie, 2019; Eberl et al., 2015; Vivier et al., 2018). In response to three classical activators, alarmin cytokines IL-33, IL-25 and TSLP, ILC2 cells can promptly produce huge amounts of type 2 effector cytokines such as IL-5 and IL-13, which drive the development of type 2 immunopathologies featured by eosinophilia, airway remodeling and mucus hypersecretion (Cayrol and Girard, 2018; Kubo, 2017; Liew et al., 2016; Molofsky et al., 2015).

In reality, environmental allergens are very complex and often contaminated with microbial or parasitic products such as lipopolysaccharide (LPS), also known as endotoxin, or chitin, so called pathogen-associated molecular patterns (PAMPs). PAMPs could stimulate innate immune cells through binding and activating their corresponding pattern recognition receptors (PRRs). Besides sensing PAMPs, PRRs can also recognize self-ligands, namely, DAMPs (damage-associated molecular patterns) (Akira et al., 2006; Brubaker et al., 2015; Man et al., 2016; Wu and Chen, 2014). Toll-like receptors (TLRs) are the first class of PRRs. All TLRs contain extracellular leucine-rich repeats (LRRs) for ligand bindings and an intracellular Toll-interleukin-1 receptor (TIR) domain that recruits MyD88 and other adaptor proteins to activate signaling cascades, which culminate in the production of various pro-or anti-inflammatory cytokines. To date, 10 TLRs (TLR1–10) have been identified in humans and 12 (TLR1–9, TLR11–13) in mice (Roach et al., 2005). TLR2 forms a heterodimer with TLR1, TLR6 or TLR10 to detect microbial lipopeptides and peptidoglycans. TLR4 detects bacterial LPS while TLR5 recognizes bacterial flagellin. The nucleic acid sensing TLRs include TLR3, TLR7/8, TLR9 and TLR13, which detect double-stranded RNA (e.g. poly[I:C]), R848 or single-stranded RNA (ssRNA), unmethylated CpG DNA (e.g. CpG-A), bacterial 23S rRNA and its derivative, a 13-nt sequence, ISR23, respectively (Li and Chen, 2012).

The immune activation of allergen-associated PAMPs is believed to be critically involved in the initiation of allergic type 2 inflammation (Locksley, 2010; Palm et al., 2012; Stewart and Thompson, 1996; Van Dyken et al., 2011; Van Dyken et al., 2017; Van Dyken et al., 2014). Recent studies have also demonstrated that microbial ligands can activate the corresponding innate immune receptors to protect against eosinophilic airway diseases through inhibition of ILC2 function(Maazi et al., 2018; Sabatel et al., 2017; Thio et al., 2019). TLR4 was previously shown to play a role in regulating type 2 immune responses in mouse studies, however, the underlying mechanism of action remains poorly understood (Deckers et al., 2017; Eisenbarth et al., 2002; Hammad et al., 2009; Hammad and Lambrecht, 2015; McAlees et al., 2015; Trompette et al., 2009). Although the role of pattern-recognition receptors (PRRs) in myeloid cells like monocytes and DCs is well established, it remains unclear whether PRRs such as TLRs can directly function in ILC2 cells.

In the course of investigating the ability of ILC2 cells to respond to various TLR ligands, we found that only LPS was able to robustly trigger human, but not murine ILC2 cells, to proliferate and produce type 2 effector cytokines including IL-4, IL-5, and IL-13. Further, LPS treatment greatly enhanced CD40 ligand (CD154) expression on ILC2 cells to promote the proliferation and antibody production of human B cells. Adoptive cell transfer experiments using humanized mouse models showed that LPS can activate human ILC2 cell in vivo in mouse lungs. Moreover, RNA-seq data revealed that LPS induced a large set of genes overlapped with those induced by IL-33. Collectively, these findings support a non-classical mode of activating human ILC2 cells by the LPS-TLR4 signaling axis.

## Results

### LPS potently activates the proliferation and cytokine production of human, but not murine ILC2 cells

ILC2 cells are known to be primarily activated by alarmin cytokines IL-33, TSLP or IL-25. However, it remains elusive whether ILC2 cells are capable of directly sensing microbial products through PRRs such as TLRs. To address this issue, various TLR agonists were initially used to examine their abilities to activate in vitro-cultured human (CD45+Lin-CRTH2+CD127+) or murine (CD45+Lin-T1/ST2+) ILC2 cells, which were originally isolated from human blood or murine lung samples by FACS sorting (**Fig. S1**). The growth and cytokine production of ILC2 cells were analyzed on day 3 (**Fig. S2**) and 5 (all experiments if not indicated) with techniques including microscopic examinations, FACS and ELISA, respectively. Among a number of TLR ligands tested, LPS (*E. coli* 0127:B8) was identified to potently stimulate human, but not murine, ILC2 cells to proliferate (**Figs. 1A & B and 1D & E**), and produced massive amounts of type 2 effector cytokines IL-5 and IL-13 (**Figs. 1C and 1F**). Pam3CSK4, a TLR2 agonist, was very toxic to human ILC2 cells, but seemed to be stimulatory to murine ILC2 cells (**Figs. 1D, E & F**). Interestingly, at a proper concentration range, Pam3CSK4 stimulated both human (0.01-0.1 μg/ml) and murine (1-10 μg/ml) ILC2 cells (**Fig. S3**), which is consistent with previous reports (Crellin et al., 2010; Ishii et al., 2019). In addition, LPS from three Gram-negative bacteria including *E. coli* 055:B5, *P. aeruginosa* and *S. typhimurium* was also found to strongly activate human ILC2 cells (**Fig. S4**). LPS activated human ILC2 cells in a concentration-dependent manner (**Fig. 2**) and promoted their survivals even at the highest concentration (100 μg/ml) (**Fig. S5**). We also carefully examined the effects of LPS at different concentrations on murine ILC2 cells and found that LPS at 10 or 100 μg/ml did have some weak stimulating effects on murine ILC2 cells **(Fig. S6)**. Taken together, these data suggested that at proper concentrations, LPS has the ability to directly activate human, but not murine ILC2 cells, to proliferate and induce the secretion of type 2 effector cytokines IL-5 and IL-13.

**Figure 1.**
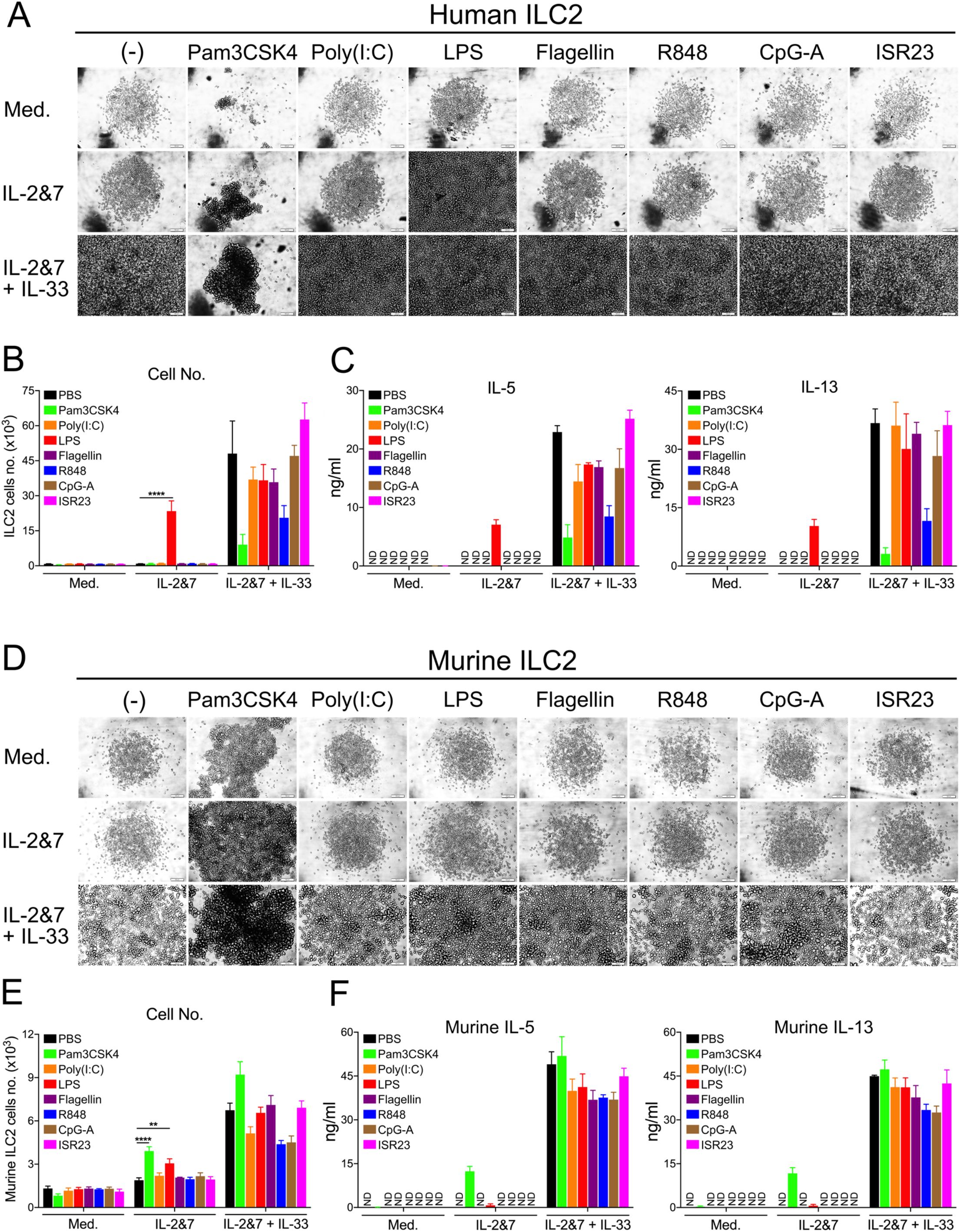
LPS strongly stimulated the growth and cytokine production of human, but not murine ILC2. **A.** Light microscopic images showing the growth of ILC2 cells. FACS sorted human ILC2 cells were treated with various TLRs as indicated in a 96-well round bottom plate for 5 days. Each image represented one well in which 1,000 cells were initially seeded. Scale bar, 100 μm. **B.** FACS showing the number of human ILC2 cells treated with various TLRs as indicated. **C**. ELISA measuring the production of IL-5 and IL-13 by human ILC2 cells treated with various TLR ligands as indicated. **D.** Light microscopic images showing the growth of ILC2 cells. FACS sorted murine ILC2 cells were treated with various TLR ligands as indicated in a 96-well round-bottom plate for 5 days. Each image represented one well in which 1,000 cells were initially seeded. Scale bar, 100 μm. **E.** FACS showing the number of murine ILC2 cells treated with various TLRs as indicated. **F**. ELISA measuring the production of IL-5 and IL-13 by murine ILC2 cells treated with various TLRs as indicated. (P value <0.05 was considered statistically significant, two-way ANOVA, ** p < 0.01, **** p < 0.0001).

**Figure 2.**
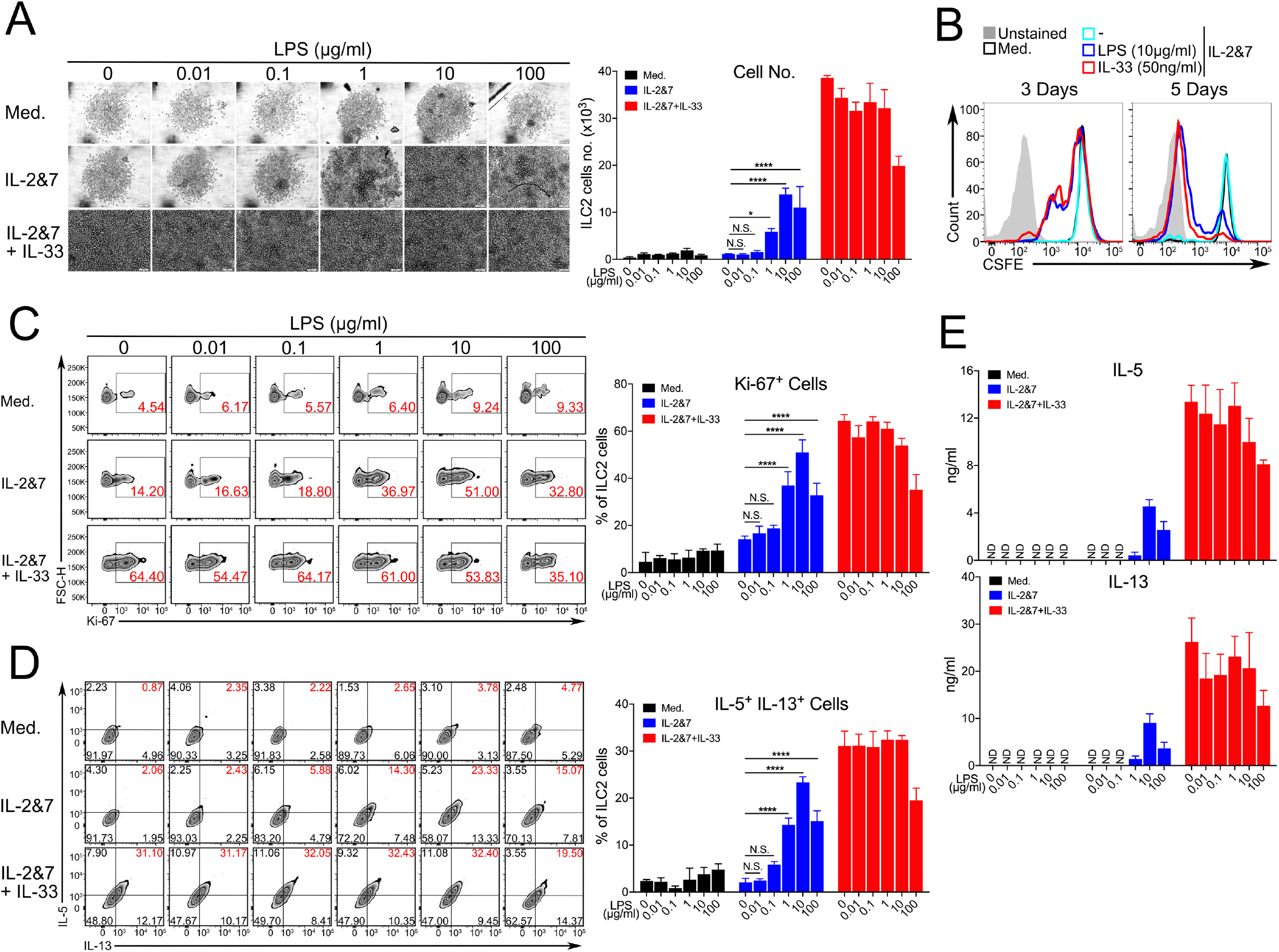
LPS stimulated human ILC2 cells in a dose-dependent manner. **A.** Light microscopic images showing the growth of ILC2 cells. FACS sorted human ILC2 cells were treated with the increased dose of LPS as indicated. Each image represented one well in which 1,000 cells were initially seeded in a 96-well round-bottom plate. Scale bar, 100 μm (left). FACS showing the number of LPS-treated human ILC2 cells (right). **B.** CSFE labelling for tacking the proliferation of human ILC2 cells treated with LPS or IL-33 for 3 or 5 days. **C.** The proliferation of LPS-treated human ILC2 cells were analyzed with Ki-67 staining. The representative FACS gating images and quantification of Ki-67 positive cells were shown. The result is a representative of two independent experiments. **D**. The percentage of IL5+IL13+-double positive human ILC2 cells treated with the increased dose of LPS were analyzed by the intracellular staining. (P value <0.05 was considered statistically significant, two-way ANOVA, **** p < 0.0001). **E.** ELISA measuring the production of IL-5 and IL-13 by human ILC2 cells treated with the increased dose of LPS.

### LPS activates human ILC2 cells via TLR4 receptor without the involvement of IL-33-ST2 pathway

Next, we investigated the involvement of TLR4- and ST2-mediated pathways in LPS-triggered signaling in human ILC2 cells. Although it was reported that human ILC2 cells expressed TLR4 mRNA and responded to a Mix of TLR-ligands (Maggi et al., 2017), it remains to be determined whether LPS alone can act on TLR4 signaling in human ILC2 cells. To address this question, we first analyzed the protein expression level of TLR4 and its co-receptor CD14 in ILC2 cells derived from 4 healthy donors. FACS staining showed that TLR4, but not CD14, was detected on the surfaces of unstimulated ILC2 cells from all 4 donors (**Fig. 3A**). Functionally, LPS strongly enhanced the growth of ILC2 cells derived from all 4 donors at various degrees. Notably, LPS could even significantly increase the effects of IL-33 on the proliferation of ILC2 cells derived from 3 donors (**Figs. 3B & C**). LPS alone triggered the cytokine production of IL-5 and IL-13 by ILC2 cells derived from all 4 donors (**Fig. 3D**). Chemically, LPS contains three parts: lipid A, O-antigen and core oligosaccharide joined by a covalent bond. Lipid A domain is responsible for the toxicity of Gram (-) bacteria and binding to TLR4. To determine a specific role of TLR4 receptor in LPS-mediated activating human ILC2 cells, we used a chemically synthesized TLR4 agonist a lipid A analog (CRX-527) (Stover et al., 2004) and an natural antagonist, LPS-RS, to stimulate or block TLR4-specific signaling, respectively. Given that the CRX-527 solvent DMSO at higher concentrations is toxic to cells, DMSO alone at different concentrations was included as a control to rule out its compound effects on ILC2 cells (**Fig. 4A**). CRX-527 at 0.1 and 1 μg/ml strongly promoted the growth (**Fig. 4A & B**) and the production of IL-5 and IL-13 (**Fig. 4C**). Due to the toxic effects of DMSO, CRX-527 at higher concentrations 10 or 100 μg/ml completely lost its stimulating effects. Next, we asked whether the LPS activity could be competitively blocked LPS-RS, a lipopolysaccharide isolated from the photosynthetic bacterium, *Rhodobacter sphaeroides*, which is known to be unable to induce TLR4 signaling (Coats et al., 2005). Remarkably, LPS-RS at 50 μg/ml or above almost completely inhibited the biological effects of LPS. It is worth to mention that both LPS and LPS-RS were dissolved in water (**Fig. 4D, E & F)**. To further address the possible involvement of IL-33-ST2 pathway, we performed a competitive assay by adding increased amounts of the recombinant protein IL1RL1 into ILC2 culture in order to sequester free IL-33. It turned out that whereas IL1RL1 at 1 and 10 μg/ml significantly blocked the effects of IL-33, it failed to affect the activity of LPS on human ILC2 cells (**Fig 5A, B & C**). These results indicate that LPS activates human ILC2 cells via TLR4 receptor without the involvement of IL-33-ST2 pathway.

**Figure 3.**
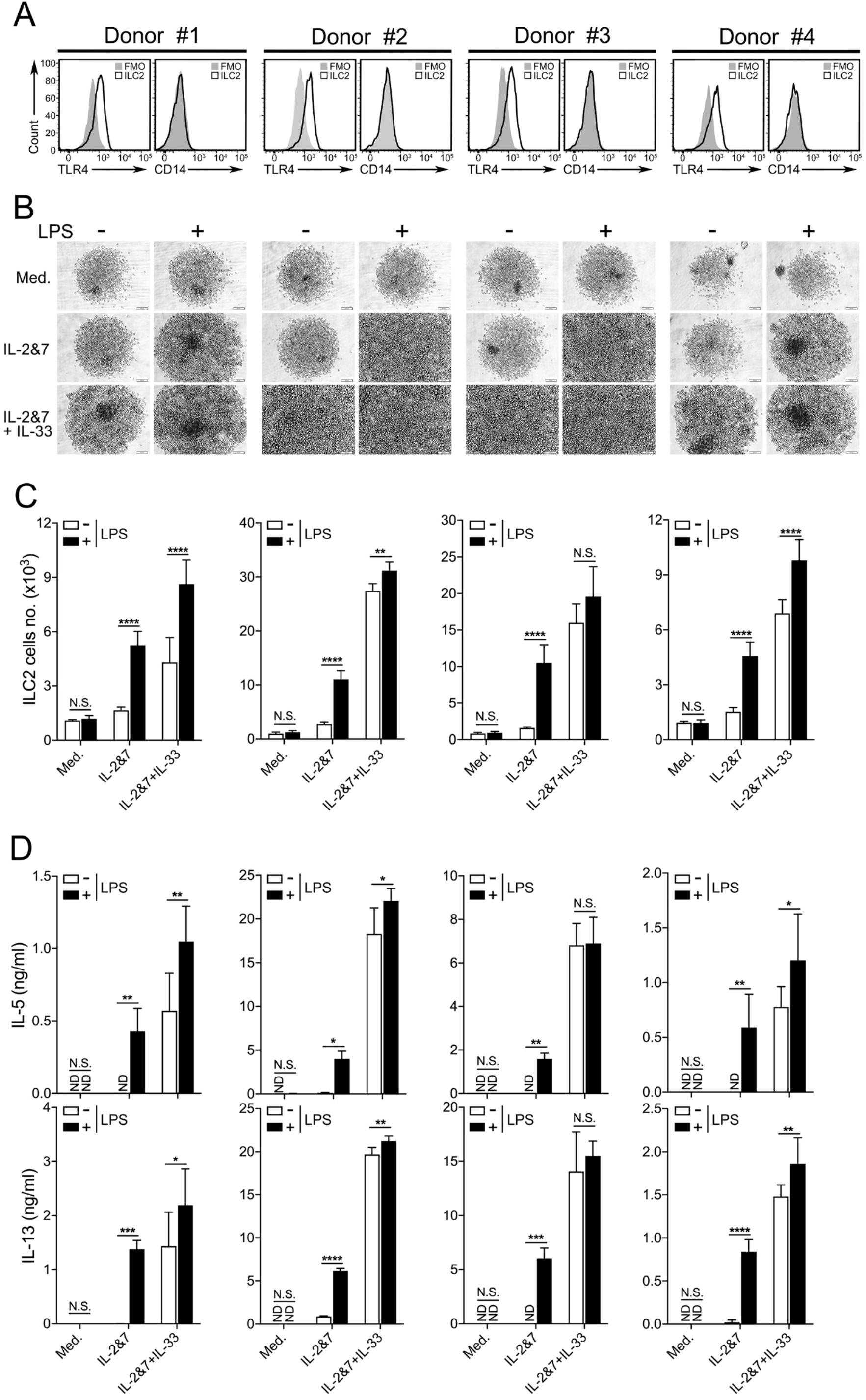
TLR4 expression and the LPS responsiveness of human ILC2 cells from four healthy donors. **A.** FACS staining of TLR4 or CD14 proteins on the surface of human ILC2 cells. FMO (Fluorescence Minus One control). **B.** Light microscopic images showing the growth of ILC2 cells derived from 4 individual donors were activated by LPS or IL-33. Each image represented one well in which 1,000 cells were initially seeded. Scale bar, 100 μm. **C.** FACS showing the number of human ILC2 cells derived from 4 individual donors were activated by LPS. **D.** ELISA measuring the production of IL-5 and IL-13 by human ILC2 cells treated with LPS. (P value≥0.05 was considered statistically insignificant, N.S., two-way ANOVA, * p < 0.05, ** p < 0.01, *** p < 0.001, **** p < 0.0001).

**Figure 4.**
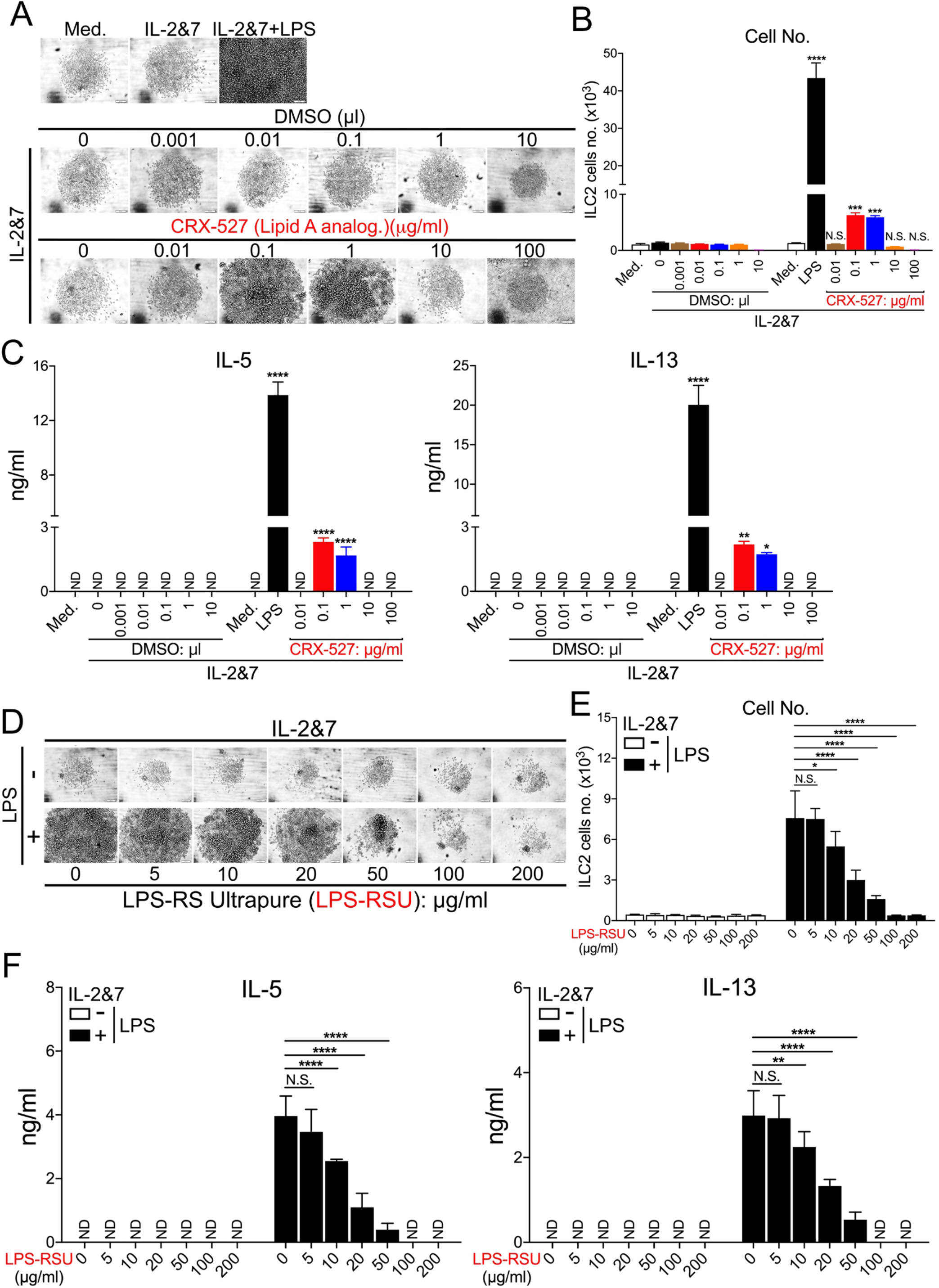
Stimulation and suppression of human ILC2 cells by the synthetic agonist or antagonist of LPS. **A.** Light microscopic images showing the growth of ILC2 cells treated with CRX-527, lipid A analog or its solvent DMSO. FACS sorted human ILC2 cells were treated with the increased dose of CRX-527 as indicated. Each image represented one well in which 1,000 cells were initially seeded. Scale bar, 100 μm. **B.** FACS showing the number of human ILC2 cells treated with either CRX-527 or DMSO. **C.** ELISA measuring the production of IL-5 and IL-13 by human ILC2 cells treated with either CRX527 or DMSO. **D.** Light microscopic images showed that the LPS-stimulated growth of ILC2 cells were inhibited by the increased dose of a specific TLR4 antagonist, LPS-RS. Each image represented one well in which 1,000 cells were initially seeded. Scale bar, 100 μm. **E.** FACS showing the number of human ILC2 cells treated with LPS-RS. **F.** ELISA measuring the production of IL-5 and IL-13 by human ILC2 cells treated with the increased doses of LPS-RS. (* P value <0.05 was considered statistically significant, unpaired t-test, ** p < 0.01, *** p < 0.001, **** p < 0.0001).

**Figure 5.**
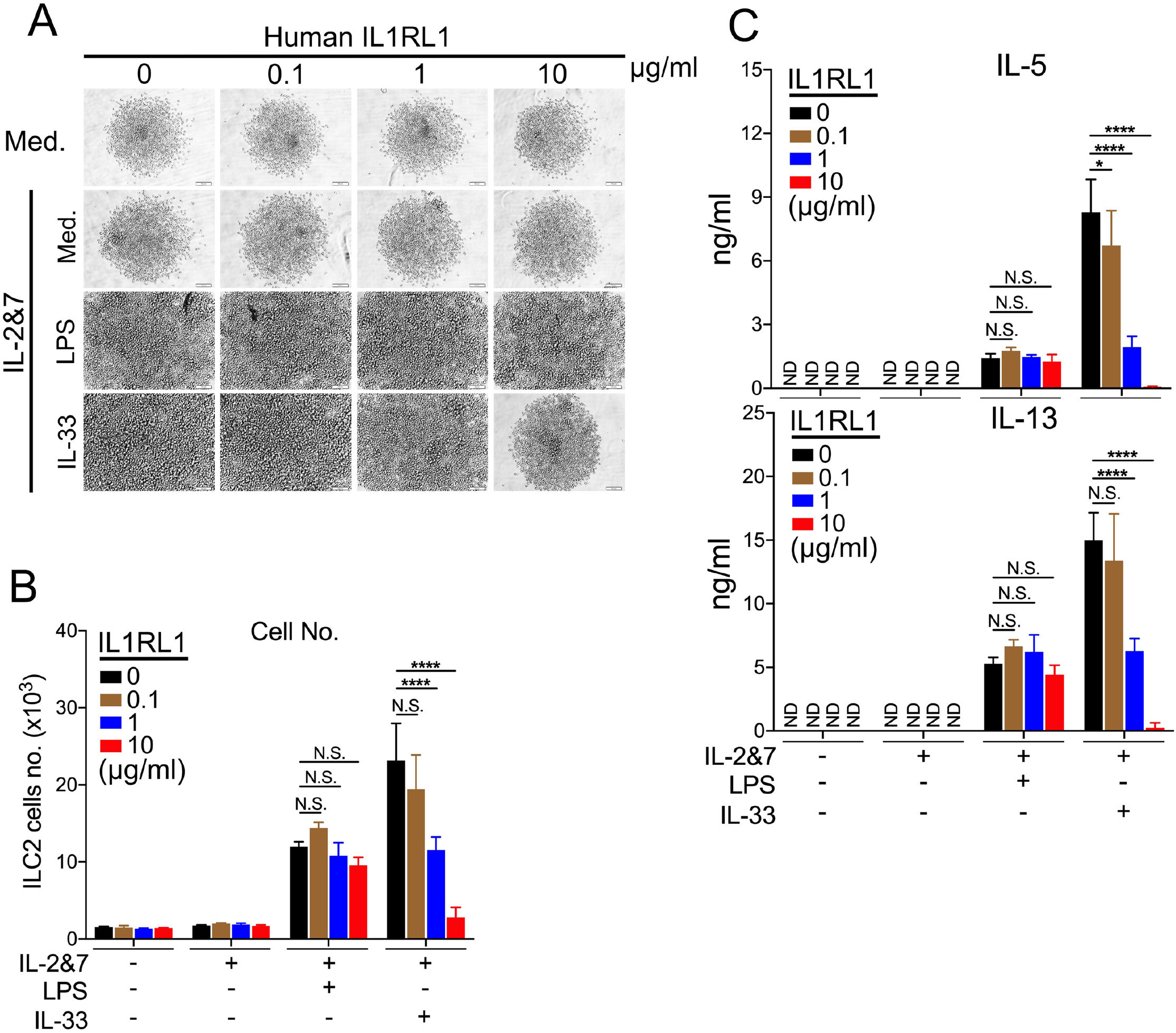
LPS-activated human ILC2 cells are insensitive to the blocking effects of recombinant protein IL1RL1, the IL-33 receptor. **A.** Light microscopic images showing the growth of ILC2 cells treated with the increased dose of recombinant protein IL1RL1. FACS sorted human ILC2 cells were activated with either LPS or IL-33 as indicated. Each image represented one well in which 1,000 cells were initially seeded. Scale bar, 100μm. **B.** FACS showing the number of human ILC2 cells treated with the increased dose of recombinant protein IL1RL1. **C.** ELISA measuring the production of IL-5 and IL-13 by human ILC2 cells treated with the increased dose of recombinant protein IL1RL1. (P value <0.05 was considered statistically significant, two-way ANOVA, **** p < 0.0001).

### RNA-seq analysis reveals that LPS-induced genes overlap significantly with those induced by IL-33 in human ILC2 cells

We wanted to obtain a global view on the LPS-activated human ILC2 cells at the transcriptional level. In order to find the best timepoint to collect samples for the RNA-seq analysis, we examined the kinetics of selected genes (IL-4, IL-5, IL-13 and TNFα) induced by LPS in comparison to an IL-33 control by RT-qPCR. We found that the peak level of these genes appeared at 6 hours in both treatments (**Fig. S7**). Therefore, we chose the 6h samples to perform an RNA sequencing analysis (**Fig. 6**). We first compared the relative mRNA expression level of all human TLRs and found that TLR4 was the highest expressed gene in non-treated human ILC2 cells, suggesting the importance of TLR4 signaling in this cell type (**Fig. 6A**). Next, we analyzed the differential expression of all genes. In a volcano-plot analysis, 213 genes were up-regulated whereas 52 genes were down-regulated compared with the control, which can be also seen in a heatmap analysis (**Fig. 6B & C**). More interestingly, further cytokine analysis found that many type 2 effector cytokines such as IL-5, IL13 and CSF2 were induced and highlighted in red (**Fig. 6D**). To further define the functional connections of the DFGs genes, the KEGG (Kyoto Encyclopedia of Genes and Genomes) pathway enrichment and GOBP (Gene Ontology Biological Process) analyses were performed. We discovered that genes from the LPS-treated sample were significantly enriched in biological processes highly related to immune modulations by cytokines such as TNF and IL-17 (**Figs. 6E & S8**). By a direct comparison using gene set enrichment analysis (GSEA), IL-33-upregulated gene set was identified to be statistically significant concordance with those induced by LPS (**Fig. 6F**), sharing more than 50% genes (141 genes) (**Fig. 6G**). Taken together, these results reveal that LPS-induced genes overlap significantly with the gene signature induced by IL-33 in human ILC2 cells.

**Figure 6.**
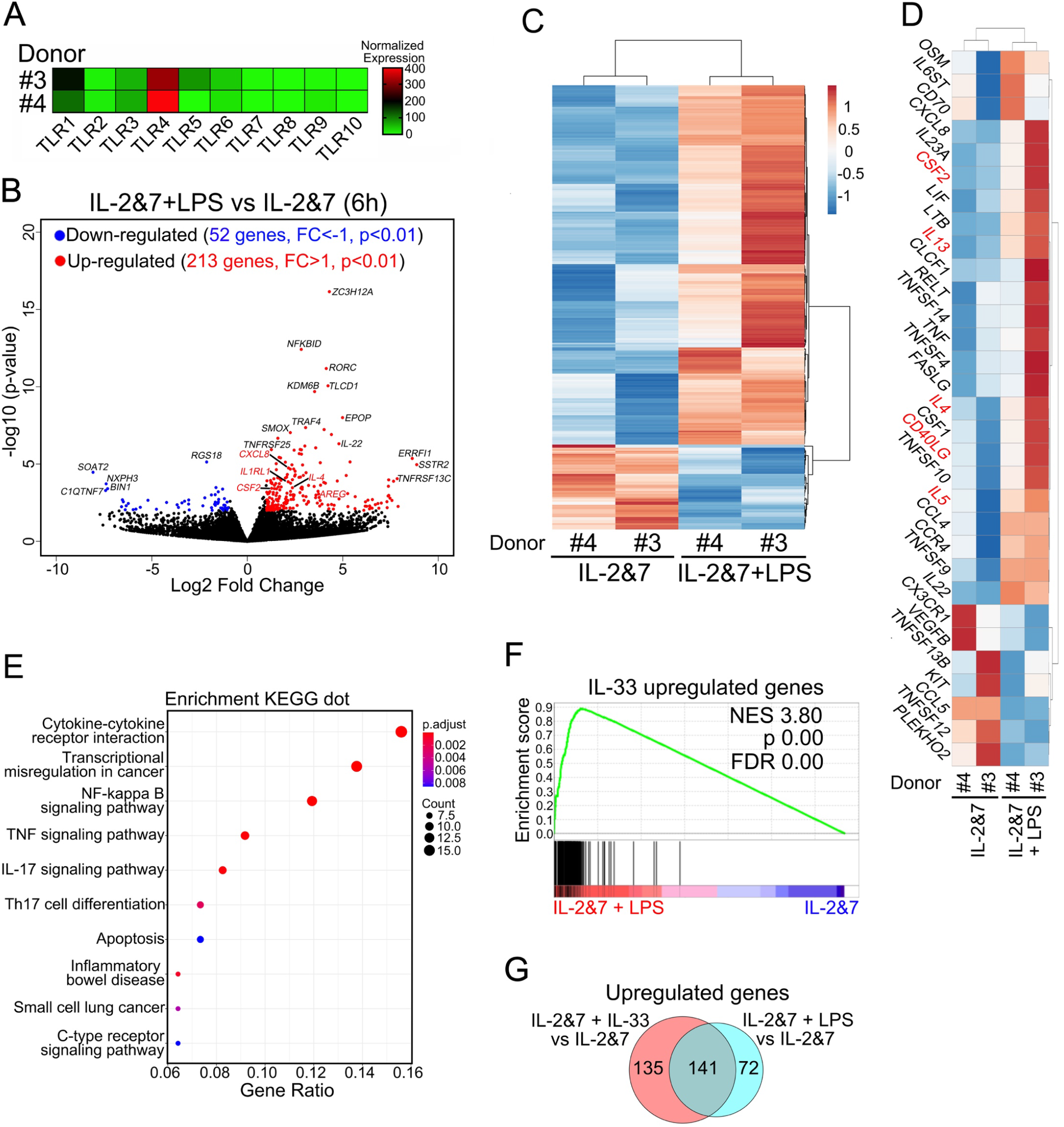
RNA-seq analysis reveals that LPS upregulated genes significantly overlapped with those upregulated by IL-33 in human ILC2 cells. **A.** Heatmap showing the normalized mRNA expression level of all 10 human TLRs in human ILC2 cells from two donors. **B.** Volcano plot showing the up- and downregulated genes in human ILC2 cells 6 hours after LPS treatment. **C.** Heatmap showing the differentially expressed genes (DEGs) in LPS-activated human ILC2 cells. **D.** Heatmap showing the selected cytokines genes in LPS activated human ILC2 cells, in which genes related to type 2 immunity are highlighted in red. **E.** Dotplot of enrichment analysis with Kyoto Encyclopedia of Genes and Genomes (KEGG). **F.** GSEA analysis of DEGs upregulated by LPS and IL-33 in human ILC2 cells. **G.** Venn diagram depicting the overlapping and non-overlapping genes upregulated in human ILC2 cells stimulated with either IL-2&7 plus LPS or IL-2&7 plus IL-33, as compared to those treated with IL-2&7.

### NF-κB and JAK-STAT pathways are critical for LPS to activate human ILC2 cells

Given that the engagement of TLRs often triggers the common signaling cascades leading to the activation of a number of transcription pathways (Kaisho and Akira, 2006), we wanted to know which pathway is required for LPS to activate human ILC2 cells. To this end, we performed additional RNA-seq analysis and found that the NF-κB and JAK-STAT signaling pathways were strongly associated with the function of LPS-activated human ILC2 cells **(Fig. 7A).** Next, we wanted to determine whether the NF-κB or JAK inhibitors would affect the growth and cytokine production of human ILC2 cells treated with LPS. As expected, NF-κB inhibition with Bay 11-7082 at the concentration between 1 to 100 μM completely blocked the LPS-mediated stimulating effects. A pan-Jak inhibitor also blocked all responses to LPS (**Figs. 7B, C & D**). Similarly, these two inhibitors also blocked IL-33-mediated effect on ILC2 cells (**Figs. 7E, F & G**). These data indicate that the NF-κB- and JAK-dependent pathways contribute significantly to LPS - mediated activation of human ILC2 cells.

**Figure 7.**
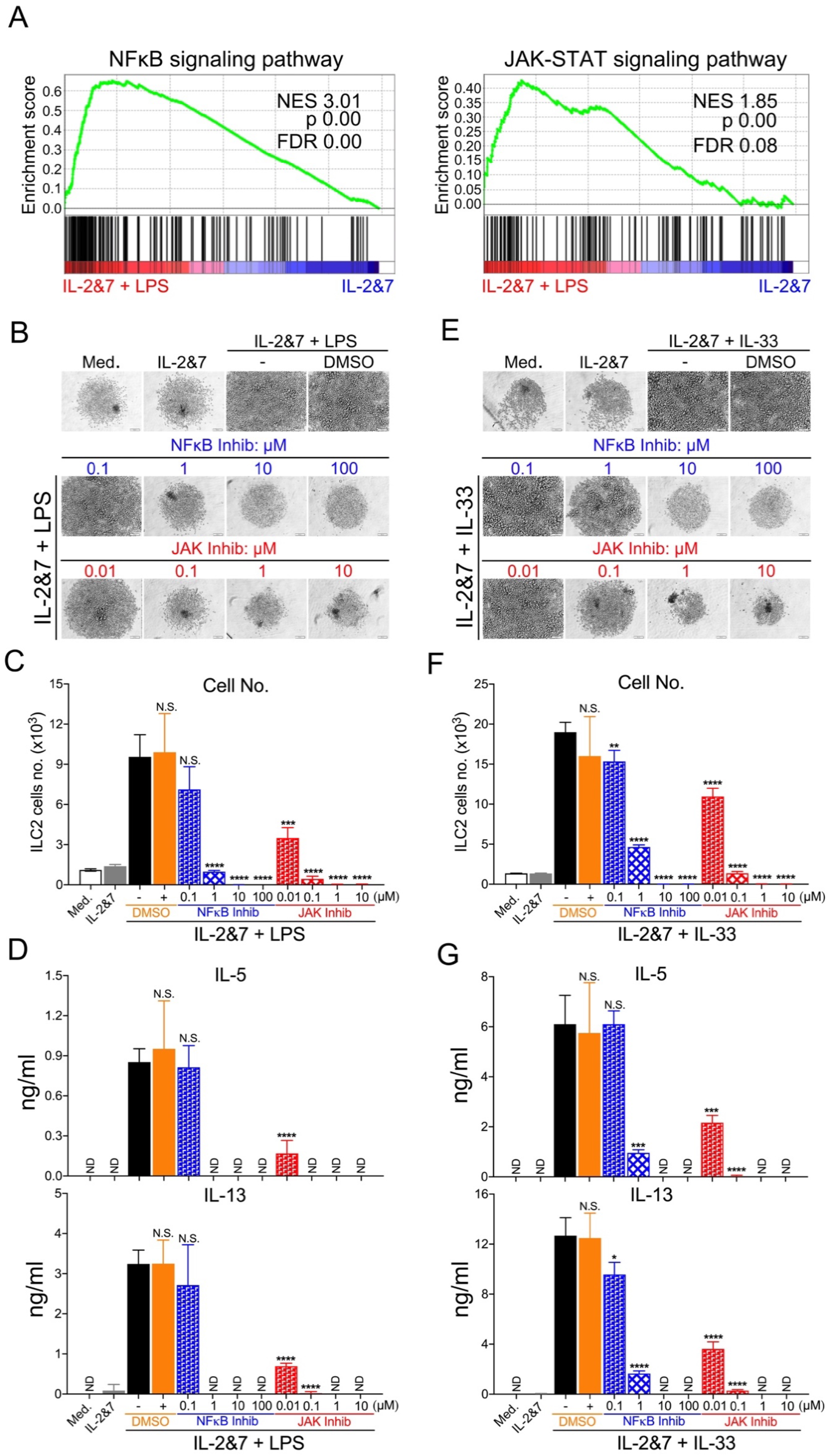
NF-κB and JAK pathways contribute to the proliferation and cytokine production of LPS- and IL-33-stimulated human ILC2 cells. **A.** Venn diagram showing the DEGs in the NF-κB and JAKSTAT pathways in human ILC2 cells stimulated with IL-2&7 plus LPS, as compared to those treated with IL-2&7 **B.** Light microscopic images showing the growth of LPS-activated ILC2 cells in the presence of the increased concentration of NF κB- and JAK-inhibitors, or DMSO. Each image represented one well in which 1,000 cells were initially seeded. Scale bar, 100μm. **C.** FACS showing the number of LPS-activated ILC2 cells in the presence of the increased concentration of NF-κB- and JAK-inhibitors, or DMSO. **D.** ELISA measuring the production of IL-5 and IL-13 by human ILC2 cells treated with LPS in the presence of the increased concentration of NF-κB- and JAK-inhibitors, or DMSO. **E.** Light microscopic images showing the growth of IL-33-activated ILC2 cells in the presence of the increased concentration of NF-κBand JAK-inhibitors, or DMSO. Each image represented one well in which 1,000 cells were initially seeded. Scale bar, 100μm. **F.** FACS showing the number of IL-33-activated ILC2 cells in the presence of the increased concentration of NF-κB- and JAK-inhibitors, or DMSO. **G.** ELISA measuring the production of IL-5 and IL-13 by human ILC2 cells treated with IL-33 in the presence of the increased concentration of NF-κB- and JAK-inhibitors, or DMSO. (P value?0.05 was considered statistically insignificant, N.S., unpaired t-test, * p < 0.05, ** p < 0.01, *** p < 0.001, **** p < 0.0001).

### LPS-activated ILC2 cells promote the proliferation and antibody production of B cells

It has been recently shown that ILC2 can interact with other cell types such as DCs, Th2 or B cells to bridge the innate and adaptive immunity (Drake et al., 2016; Gurram and Zhu, 2019; Halim et al., 2016; Maggi et al., 2017; Moon et al., 2004). We wondered whether LPS-activated ILC2 cells could activate naïve B cells. Notably, it is generally believed that LPS is unable to directly activate human B cell because its receptor TLR4 expression of human B cell is very low (Bourke et al., 2003; Ganley-Leal et al., 2010). Since we have shown above that LPS-activated human ILC2s increased the mRNA or protein expression of a few B cell regulatory factors including CD40LG (CD154), IL-5 and IL-13 (**Figs. 3D & 6D**) (Defrance et al., 1994; Moens and Tangye, 2014; Moon et al., 2004), we then asked whether the LPS-activated human ILC2s cells could activate naïve B lymphocytes by producing other B cell stimulating factors at the protein level such as CD154 and IL-4, which are required for the antibody class switching. Indeed, the CD154 expression and the secretion of IL-4 by ILC2 cells from all 4 donor was greatly induced after the treatment with either LPS or IL-33 (**Figs. 8A & B**). To further prove that LPS-treated human ILC2s were able to induce the antibody production of human B cells, we purified human B cells (CD19^+^ cells). As illustrated in **Fig. 8C**, ILC2s cells were cocultured with B cells at the 1:1 ratio in the absence or presence of LPS, respectively. After 5 days, the number of B cells and levels of IgM, IgG, IgA and IgE present in culture supernatants were measured. As shown in **Figs. 8D & E**, LPS-stimulated ILC2 activated B cells to proliferate and produce high levels of antibodies including IgM, IgG, IgA and IgE. Unstimulated ILC2 cells failed in inducing the growth and immunoglobulin production of B cells. Collectively, these results suggest that LPS-TLR4 signaling in human ILC2 is sufficient to activate the proliferation and antibody production of human B cells.

**Figure 8.**
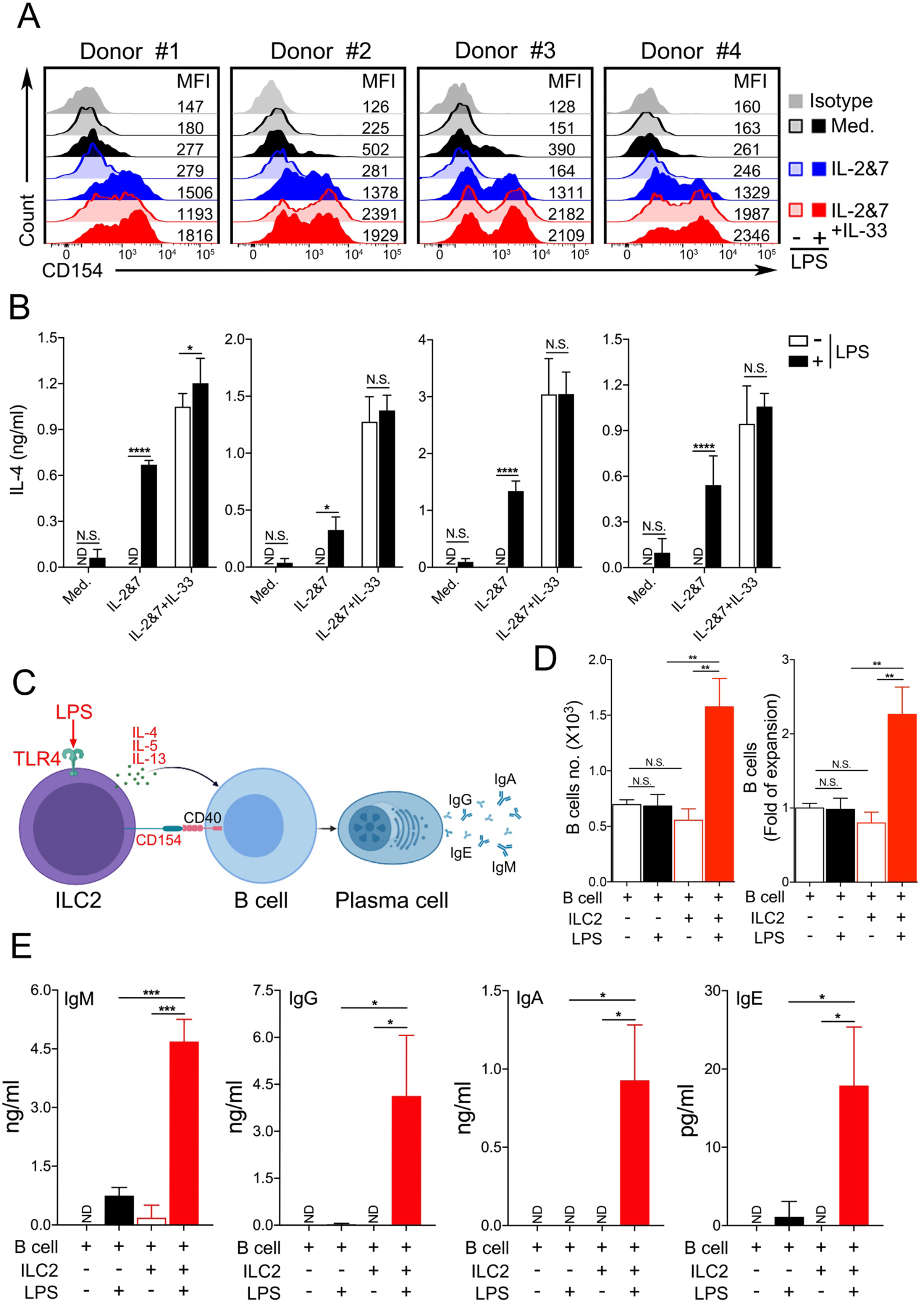
LPS-activated human ILC2 cells enhanced the proliferation and antibody production of B cells. **A.** FACS staining of CD154 proteins on the surface of human ILC2 cells from 4 individual donors treated with or without LPS as indicated. **B.** ELISA measuring the production of IL-4 by human ILC2 cells from 4 individual donors treated with or without LPS as indicated. **C.** A schematic diagram describing a working model for the interplay between LPS-activated ILC2 and B cells. **D.** FACS showing the increased number of B cells co-cultured with human ILC2 cells in the presence or absence of LPS. **E.** ELISA measuring the levels of antibodies (IgM, IgG, IgA and IgE) in supernatants of the co-cultured B and ILC2 cells in the presence or absence of LPS. (P value≥0.05 was considered statistically insignificant, N.S., unpaired t-test, * p < 0.05, ** p < 0.01, *** p < 0.001, **** p < 0.0001).

### LPS induces human ILC2 cells-mediated eosinophilic lung inflammation in vivo

Since we have shown that LPS is a potent ILC2 activator in vitro, we next moved on to evaluate whether LPS was able to elicit human ILC2 cells-mediated eosinophilia in vivo. To this end, we developed two humanized mouse models, in which NSG or Rag2^−/−^γc^−/−^ mice were reconstituted with human ILC2s as illustrated in **Fig. 9A**. Similar models have recently been reported with the IL-33 administration (Crellin et al., 2010; Hurrell et al., 2019). Due to a high level of protein sequence homology, human IL-5 could function as murine IL-5 and was previously used to activate murine eosinophils (Ingley et al., 1991; Tavernier et al., 1995). Consistent with our in vitro data, human ILC2 cells in mouse lung responded to LPS stimulations, which led to increased number of murine lung eosinophils in both NSG or Rag2^−/−^γc^−/−^ mice adoptively transferred with human ILC2 cells (**Fig. 9B**). Moreover, we also observed that the growth and type 2 cytokine production of human ILC2 cells were greatly increased by LPS treatment shown by the FACS cell number and intracellular cytokine staining in these two mouse strains (**Figs. 9C & D**). These data indicate that LPS can function in vivo to initiate human ILC2 cells-mediated eosinophilic lung inflammation.

**Figure 9.**
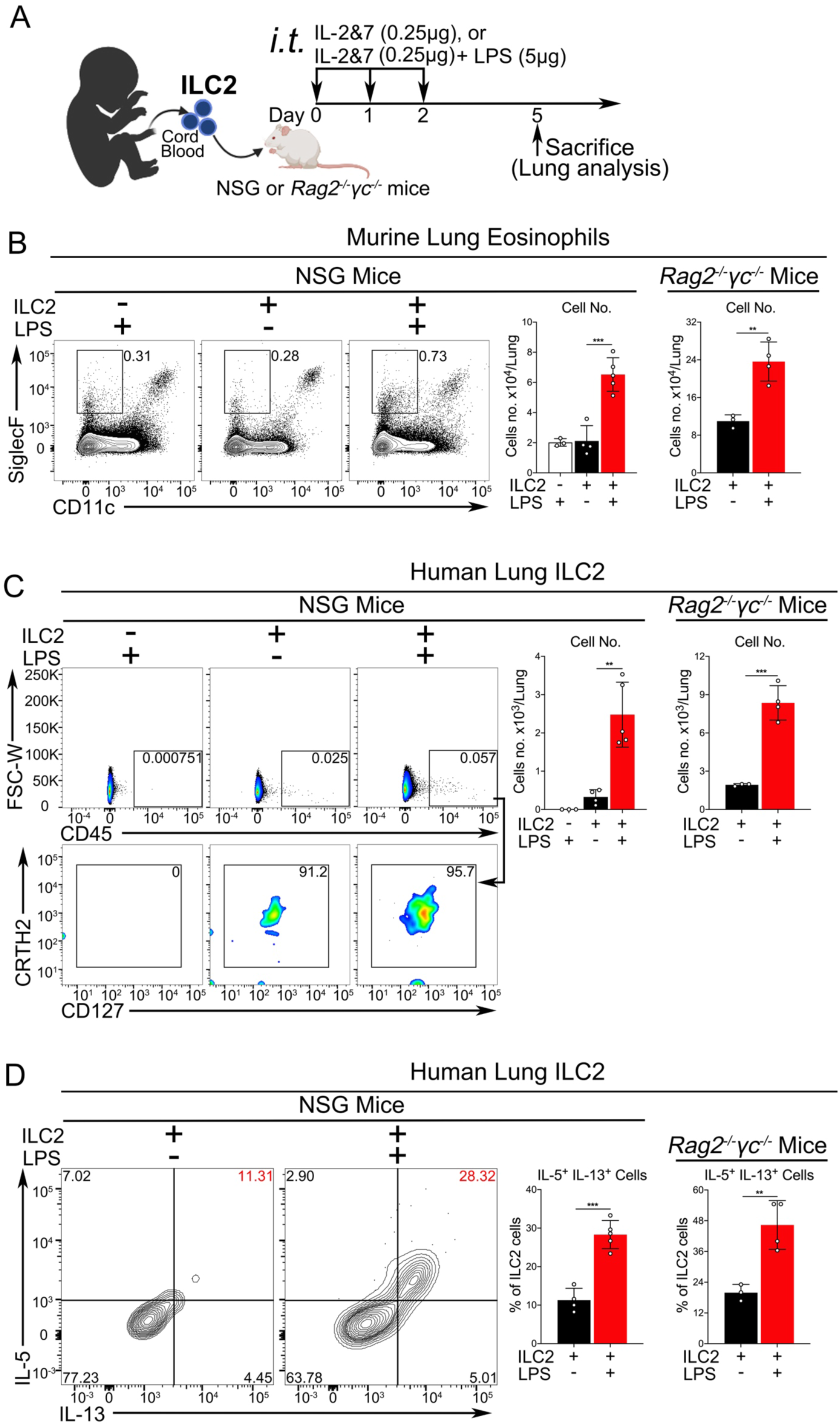
LPS could induce human ILC2 cells-mediated eosinophilic lung inflammation in humanized mouse models. **A.** A experimental protocol for studying the activation of human ILC2 cells by LPS in twohumanized mouse models. **B.** FACS showing the number of murine lung eosinophils in NSG or Rag2^-/-^*γ*c^-/-^ mice, which were reconstituted with human ILC2 and treated with or without LPS. **C.** Same as **B**, FACSshowing the number of human ILC2 cells in NSG or Rag2^-/-^*γ*c^-/-^ mice. **D.** Same as **B**, the percentage ofIL5+IL13+-double positive human ILC2 cells in NSG or Rag2^-/-^*γ*c^-/-^ mice were analyzed by the intracellularstaining. (P value≥0.05 was considered statistically insignificant, N.S., unpaired t-test, ** p < 0.01, *** p < 0.001,).

## Discussion

In this study, we have discovered a novel function for LPS-TLR4 signaling in activating ILC2 cells. We found that LPS extracted from multiple species of Gram-negative bacteria can potently stimulate human, but not murine ILC2 cells, to proliferate and produce massive amounts of type 2 effector cytokines IL-4, IL-5, and IL-13. This activation of ILC2 cells by the LPS-TLR4 signaling axis does not require the classical IL-33-ST2 pathway but rather requires the NF-kB and JAK pathways. More interestingly, LPS dramatically increased CD40 ligand (CD154) protein level on ILC2 cells to drive the proliferation and antibody production of human B cells. Further, LPS can work in vivo to activate human ILC2 cell in mouse lungs in a humanized mouse model. Mechanistically, RNA-seq data strikingly revealed that LPS upregulated a large set of genes overlapped significantly with those induced by IL-33. To our best knowledge, this is the first time to demonstrate that LPS-TLR4 signaling directly functions in human ILC2 cells to promote their proliferation and production of type 2 effector cytokines independent of the endogenous IL-33-ST2 pathway.

ILC2 cells are known to be activated by tissues-derived alarmin signals when exposed to environmental allergens. In recent years, accumulating evidence has also suggested that diverse receptors expressed on the surface of ILC2 cells can either enhance or repress their activation and proliferation in response to various non-classical signals, hormones, regulatory cytokines, neuropeptides and lipids (Hurrell et al., 2018). Our findings have identified that TLR4 signaling pathway plays an important role in regulating the survival, proliferation, and the production of type 2 cytokines by human ILC2 cells. Although it was previously shown that human ILC2 cells expressed the mRNA of some TLRs (TLR1, TLR4, and TLR6) and could respond to a mixture of three TLR-ligands (Maggi et al., 2017), it has remained unknown about which TLR pathway actually functions in human ILC2 cells.

Many previous epidemiological and experimental studies have suggested that endotoxin-TLR4 signaling contributes to the development of type 2 inflammatory reactions (Idzko et al., 2007; Janeway, 1989; Kool et al., 2011; Schuijs et al., 2015; Trompette et al., 2009). However, its underlying cellular and molecular mechanisms remain poorly understood. Natural allergens contain not only allergic proteins, but also various PAMPs derived from bacteria, parasites and viruses, which may stimulate innate immune sensors such as TLRs to participate in the regulation of type 2 inflammatory responses (Locksley, 2010; Palm et al., 2012; Stewart and Thompson, 1996; Van Dyken et al., 2011; Van Dyken et al., 2017; Van Dyken et al., 2014). In the current study we have demonstrated that bacterial endotoxin could directly engage TLR4 receptor expressed on the surface of ILC2 cells to robustly activate their proliferation and production of type 2 effector cytokines in culture and in vivo in a humanized mouse model. Besides its classical ligand, LPS, many endogenous ligands such as HMGB1(Tang et al., 2007), hyaluronan (Jiang et al., 2005), OxPAPC (Imai et al., 2008), surfactant protein A (Guillot et al., 2002), S100 proteins (Vogl et al., 1999), HSP72 (Chase et al., 2007; Wheeler et al., 2009) etc. have been reported to act as TLR4 agonists. It has been recently reported that the proteinase-cleavage of fibrinogen could elicits allergic responses through TLR4-mediated pathway (Millien et al., 2013). It would be interesting to evaluate whether those self TLR4 ligands are able to activate human ILC2 cell in future studies.

ILC2 cells have the ability to communicate with various cell types such as epithelial cells, dendritic cell, Th2 cells and mouse B cells to regulate the tissue-specific immunity (Gurram and Zhu, 2019; Hurrell et al., 2018; Klose and Artis, 2020). In agreement with a recent report showing human ILC2s can express CD154 and stimulate B lymphocytes through IL-25/IL-33 or a mixture of TLR ligands (Maggi et al., 2017), we identified that LPS treatment alone greatly enhanced CD40 ligand (CD154) expression on ILC2 cells and strongly induced three B cell regulatory cytokines IL-4, IL-5, and IL-13, which collectively promote the proliferation and antibody production of human B cells. However, more works are needed in order to understand the underlying mechanisms of antibody-class switching in this context and in vivo significance of these interesting findings.

In conclusion, our findings support a new mode of activating human ILC2 cells via the LPS-TLR4 signaling axis without the involvement of its classical activating pathway mediated by IL-33 receptor, IL1RL1. As to the wide availability of TLR4 ligands, which can originate from either a foreign origin such as bacterial infections or an endogenous source like DAMPs, thus, targeting TLR4 signaling pathway might be developed as a new approach for improving ILC2-mediated type 2 inflammatory conditions.

## Material and methods

### Mice

*Rag1*^*−/−*^ mice, *Rag2*^*−/−*^*γc*^*−/−*^ (*Rag2*^*tm1.1Flv*^*Il2rg*^*tm1.1Flv*^/J, 014593) mice and NSG (NOD.Cg-*Prkdc*^*scid*^*Il2rg*^*tm1Wjl*^/SzJ, 005557) mice were purchased from the Jackson Laboratory. Mice were bred and maintained under specific pathogen-free conditions in the animal facility according to the experimental protocols approved by the Institutional Animal Care and Use Committee.

### Regents

The medium used was RPMI 1640 (Sigma) containing 10% FBS (HyClone), 1% penicillin-streptomycin (Gibco), 1x GlutaMAX™-I (Gibco) and 50 μM 2-Mercaptoethanol (Sigma). PamsCSK4, Poly(I:C), Flagellin, R848 and CpG-A were purchased from InvivoGen. ISR23 was obtained from IDT (Integrated DNA Technologies). LPS from *Escherichia coli* 0127:B8, *Escherichia coli* 055:B5, *Pseudomonas aeruginosa* 10 (*P. aeruginosa*), *Salmonella enterica serotype typhimurium* (*S. typhimurium*) were all purchased from Sigma. Recombinant cytokines IL-2, IL-7 and IL-33 (human and mouse) were from PeproTech. TLR4 agonist: CRX-527 (Cat. #tlrl-crx-527) and TLR4 antagonist: LPS-RS Ultrapure (Cat.# tlrl-prslps) were obtained from InvivoGen. Human IL1RL1/ST2 Protein (isoform a, His Tag) (Cat. #13034-H08H) was purchased from Sino Biological. NF-κB inhibitor Bay 11-7802 and JAK inhibitor 1 were both from EMD Millipore. PMA and Brefeldin A were purchased from Sigma and BioLegend, respectively.

### Isolation of ILC2 cells

Human ILC2s were isolated from peripheral blood of healthy donors or umbilical cord blood samples from healthy full-term births in the Department of Obstetrics and Gynecology of UT Health San Antonio. All human samples were used in compliance with UT Health San Antonio Institutional Review Board. Peripheral or Cord Blood Mononuclear Cells (PBMCs or CBMCs) were isolated from diluted umbilical cord blood (1:2) by density gradient centrifugation using density gradient medium, Histopaque^®^ (Sigma Aldrich) and SepMateTM 50 mL tubes (STEMCELL Technologies) (Grievink et al., 2016). Cells were then washed once with dPBS-FBS buffer (dPBS, 3% fetal bovine serum, 1mM EDTA) and resuspended in dPBS-FBS. Cells were stained with antibodies against CD45 and lineage markers (CD3, CD14, CD16, CD19, CD20 and CD56), and ILC2 markers CRTH2, CD127 (all from BioLegend). Human ILC2s were sorted by the BD FACSAria cell sorter as CD45^+^Lin^-^CRTH2^+^CD127^+^ cells. The purity of sorted ILC2s was determined to be greater than 95%. Sorted human ILC2s were cultured and expanded in medium supplemented with rh-IL-2 and rh-IL-7 (all at 50 ng/mL) in 96-well round plates for 6 days before further experiments.

Murine Lung ILC2s were isolated from *Rag1*^*−/−*^ mice treated with rm-IL-33 (250 ng/mouse) for 3 consecutive days plus 2 days of resting before processing lung tissues for sorting ILC2 cells with an BD FACSAria cell sorter. The criteria for identifying ILC2 is lacking classical lineage markers (CD3∊, CD4, CD8α, CD11c, FceRIα, NK1.1, CD19, TER119, CD5, F4/80 and Gr-1), but expressing markers of CD45 and T1/ST2 (all from BioLegend). The purity of sorted ILC2s should be greater than 95%. Sorted ILC2s were cultured and expanded in medium supplemented with murine IL-2 and IL-7 (all at 10 ng/mL) in 96-well round plates for 6 days before further experiments.

### Culture and treatment of human and murine ILC2 cells

Sorted human ILC2 were cultured in the medium (200 μl) with or without rh-IL-2, rh-IL-7 and rh-IL-33 (all at 50 ng/ml) in 96-well round plates (1,000 cells/well) in a 37°C incubator with 5% CO_2_. Cells were treated with Pam3CSK4, Poly(I:C), LPS, Flagellin, R848, CpG-A and ISR23 (different concentration as indicated) for 3 or 5 days. And the human ILC2 cells were treated with TLR4 agonist or antagonist, human IL1RL1/ST2 Protein (isoform a, His Tag) (added 1 hour prior to the treatment with LPS or IL-33), NF-κB or JAK inhibitor as indicated in figure legends for 5 days. After 3 days treatment, the percentage of IL-5^+^IL13^+^ cells, the expression of Ki-67 and cell death of ILC2 cells were analyzed by flow cytometry. After 5 days treatment, the number and proliferation of ILC2 cells were analyzed by flow cytometry, and the levels of cytokines (IL-4, IL-5 and IL-13) in the supernatants were measured by ELISA.

Sorted murine lung ILC2 were cultured and treated with the same TLR ligands in 200 μl media with or without murine IL-2, IL-7 and IL-33 (all at 10ng/ml) in 96-well round plates (1,000 cells/well) in a 37°C incubator with 5% CO2. The percentage of IL-5^+^IL13^+^ cells, expression of Ki-67 on murine ILC2 cells were analyzed by flow cytometry 3 days later. The number and proliferation of ILC2 cells were analyzed by flow cytometry 5 days later, and the supernatant were collected for further detecting of IL-5 and IL-13 by ELISA.

### Flow cytometric analysis

Fc receptors were blocked with 2.4G2 hybridoma supernatant (generated in the lab). For the detection of early-stage apoptotic cells, Annexin V staining was performed according to the protocol (eBioscience). Intranuclear staining of Ki-67 was performed with the True-Nuclear™ Transcription Factor Buffer Set (BioLegend) according to the manufacturer’s instructions. For intracellular cytokine staining, human or murine ILC2 cells were cultured and treated as indicated for 3 days, then followed by incubating with Brefeldin A for 3 hours. After surface staining, cells were fixed and permeabilized with BioLegend Cytofix/Perm buffer and further stained intracellularly with anti-human IL-5 and IL-13, or anti-mouse IL-5 and IL-13, respectively. For lung single-cell suspensions, 2 × 10^6^ total live nucleated cells were stimulated in 200 μl media with Brefeldin A and PMA (phorbol 12-myristate 13-acetate) (30 ng/mL) at 37 °C for 3 hours. After surface staining, cells were fixed and permeabilized and further stained intracellularly with anti-human IL-5 and IL-13. Dead cells were stained with eFluor506 Fixable Viability Dye before fixation and permeabilization and excluded during analysis. For TLR4 and CD14 expression on human ILC2 cells, cells were stained with the surface markers anti-Human TLR4 and anti-Human CD14. For CD154 expression, human ILC2 cells were cultured and stimulated as indicated for 5 days, and followed by staining with anti-Human CD154.

### Protein quantification in cell culture supernatants

Cytokines (IL-5 and IL-13) in supernatant of human and murine ILC2 cell cultures were analyzed with ELISA kits from Invitrogen. For human cytokine IL-4 in the supernatant of human ILC2 cell cultures were measured by ELISA kit from BioLegend. All final reactions were developed with TMB substrate (Thermo scientific) and stopped by sulfuric acid (0.16M), and the OD at 450 nm was measured.

### CSFE staining

Human ILC2 cells were stained with 1 μM CSFE (CSFE cell division tracker kit, BioLegend) according to the manufacturer’s recommendations. Followed by cultured in the medium with or without rh-IL-2, rh-IL-7, rh-IL-33 (all at 50 ng/ml) or LPS (10 μg/ml) for 3 or 5 days in a 37°C incubator with 5% CO_2_. The proliferation of ILC2 cells were analyzed by flow cytometry.

### Humanized Mice

Purified human ILC2 cells were cultured with rh-IL-2 and rh-IL-7 (all at 50 ng/ml) for 6 days, and then adoptively transferred to NSG mice or *Rag*^*−/−*^*γc*^*−/−*^ mice (4 × 10^4^ cells/mouse). 4 hours after cell transfer, host mice were challenged with rh-IL-2&7 (all 250 ng/mouse) in absence or presence of LPS (5 μg/mouse) on day 0, 1, 2 as shown in **Fig. 9A**. On day 5, mice were sacrificed, lung was performed and analyzed.

### FACS analysis of lung

Mice were sacrificed at indicated times and the lung tissues were digested in 8 ml RPMI-1640 containing Liberase (50 μg/ml) and DNase I (1 μg/ml) for about 45 min at 37 °C. Cell suspensions were filtered through 70 μm cell strainers and washed once with RPMI-1640. Human ILC2 cells and mouse eosinophils in lung were labeled with antibodies as indicated, then mixed with counting beads (Spherotech) for further FACS analysis on BD Celesta cell analyzer. Flow cytometry data were analyzed using FlowJo software. The antibodies and reagents for FACS analysis are listed below: SPHERO™ AccuCount Fluorescent (Spherotech, Cat.# ACFP-70-5), Anti-Human CD45 APC-Cy7 (HI30, BioLegend), Anti-Human CRTH2 APC (BM16, BioLegend), Anti-Human CD127 PE (A019D5, BioLegend), Anti-Mouse Siglec-F PE (E50-2440, BD Bioscience), Anti-Mouse CD11c PE-Cy7 (N418, TONBO bioscience), Anti-Mouse CD45 PerCP-Cy5.5 (30-F11, BioLegend), Fixable Viability Dye eFluor 506 (Invitrogen).

### ILC2 and B cells co-culture and Immunoglobulin production

Human naive CD19^+^ B cells (>95% pure) were purified by negative selection, using the EasySep Human Naive B Cell Enrichment Kit (19254; STEMCELL Technologies), from healthy donor PBMCs, following the manufacturer’s instructions. Naïve B cells (2,000/well) were cultured in RPMI medium with rhIL-2&7 (all at 50 ng/ml) or cocultured with ILC2 cells (2,000/well) in the presence of LPS (10 μg/ml) in 96-well round plates for 5 days. Supernatants from each well were collected and further evaluated of IgG, IgM, IgA and IgE by a homemade ELISA. Except that 96-well plates were coated with F(ab’)2 anti-Ig (H+L) Ab, and captured Igs were detected by biotinylated anti-IgG, -IgA, -IgE and -IgM Abs. Followed by reaction with HRP-labeled streptavidin, development with TMB substrate (Thermo scientific) and stopped by sulfuric acid (0.16M), the OD at 450 nm was measured.

### Real-Time Quantitative PCR

Human ILC2 cells were treated with or without LPS (10 μg/ml) or rh-IL-33 (50 ng/ml) in 200 μl media with or without rh-IL-2&7 (all 50 ng/ml) for 6 or 16 hours. Total RNA was extracted with TRIzol reagent (Life Technologies) according to the manufacturer’s instructions. Reverse transcription and real-time PCR (qPCR) reactions were carried out using iScript cDNA synthesis kit and iQ SYBR Green Supermix (Bio-Rad). qPCR was performed on a Bio-Rad CFX384 Touch™ Real-Time PCR Detection System using the following human primers (Forward and Reverse, 5’→3’): GAPDH (ATGACATCAAGAAGGTGGTG; CATACCAGGAAATGAGCTTG), IL-4 (ACTTTGAACAGCCTCACAGAG; TTGGAGGCAGCAAAGATGTC), IL-5 (AGCTGCCTACGTGTATGCCA; CAGGAACAGGAATCCTCAGA), IL-13 (TGAGGAGCTGGTCAACATCA; CAGGTTGATGCTCCATACCAT), TNFα (CCTGGTATGAGCCCATCTATCTG; TAGTCGGGCCGATTGATCTC).

### RNA isolation, RNA-Seq and Bioinformatics

Purified human ILC2 cells (4×10^4^) were cultured and stimulated with LPS (10 μg/ml) or recombinant human IL-33 protein (rh-IL-33) (50 ng/ml) in the presence with rh-IL-2&7 (all at 50 ng/ml) for 6 hours. Total RNA was extracted with RNeasy Mini Kit (Qiagen) according to the manufacturer’s instructions.

After the quality of RNA samples was verified with an Agilent Bioanalyzer 2100 (Agilent), RNA was further processed using an Illumina TruSeq RNA sample prep kit v2 (Illumina). Clusters were generated using TruSeq Single-Read Cluster Gen. Kit v3-cBot-HS on an Illumina cBot Cluster Generation Station. After the quality control procedures, individual RNA-Seq libraries were pooled based on their respective 6 bp index portion of the TruSeq adapters and sequenced at 50 bp/sequence using an Illumina HiSeq 3000 sequencer. Resulting reads were checked by assurance (QA) pipeline and initial genome alignment (Alignment). After sequencing, demultiplexing with CASAVA was employed to generate a Fastq file for each sample. All sequencing reads were aligned with the (GRCh38/hg38) reference genome using HISAT2 default settings, yielding Bam files, duplicated reads were discarded by using Picard, then were processed using HTSeq-count to obtain counts for each gene. RNA expression levels were determined using GENCODE annotation. Differential expression analysis was performed using the Deseq2 package in R post-normalization based on a Benjamini-Hochberg false discovery rate (FDR)-corrected threshold for statistical significance of padj <0.05 or raw p value <0.01. Transcript read counts were transformed to ln(x+1) used to generate heatmaps in Clustvis. Volcano plots depicting log2-FoldChange and raw or adjusted p values were generated in R. Venn diagram were generated by use the VennDiagram package in R.

To investigate biological pathways, DEGs (Differential Expression Genes) were manually curated and compared to multiple public databases, including Gene Ontology (GO) and Kyoto Encyclopedia of Genes and Genomes (KEGG), for enrichment analysis. For gene set enrichment analysis (GSEA), IL-33 upregulated gene set was identified from our RNASeq data comparing the sample rh-IL-33 in the presence of rh-IL-2&7 with the sample only has rh-IL-2&7 stimulation by using the threshold for statistical significance raw p value <0.01 and log2-FoldChange >1. Samples with or without LPS in the presence of rh-IL-2&7 were directly compared to this gene set to identify statistically significant concordance in the expression using the gene set enrichment analysis (GSEA) algorithm.

### Statistical analysis

The statistical analysis was done using software GraphPad Prism 6. For comparison of two groups, P values were determined by unpaired two-tailed Student’s t test. For comparison of more than two groups, Two-Way ANOVA was performed. P values are indicated on plots and in figure legends. P value<0.05 was considered statistically significant. (P value ≥0.05 was not considered statistically significant [N.S.]. * p < 0.05, ** p < 0.01, *** p < 0.001, **** p < 0.0001).

## Acknowledgements

We thank Ms. Karla Gorena, Mr. Sebastian Montagnino for technical assistance in flow cytometry and FACS sorting. We thank Drs. Zhao Lai, Yi Zou, and Yidong Chen for RNA-seq analysis and informatics assistance. L.S. is supported by the China Scholarship Council and Hunan Provincial Innovation Foundation for Postgraduate (CX201713068). H.H.A. is supported by the Department of Clinical Laboratory Sciences, College of Applied Medical Sciences, Jouf University, Sakaka, Saudi Arabia. X.-D.L. is supported by the UT Health San Antonio School of Medicine Startup Fund and the Max and Minnei Voelcker Fund.

## Author contributions

L.S. and H.H.A. performed most experiments; L.S., H.H.A., J.W., D.P.C., Y.X., H.Z., Z.X., Y.S., N.X., X.Z., Y.L. and X.-D.L. analyzed data; L.S., H.H.A., Y.L. and X.-D.L. planned, designed research. L.S., Y.L. and X.-D.L. wrote the manuscript; All authors discussed the results and participated in writing and commenting on the manuscript.

---

**Fig. S1.**
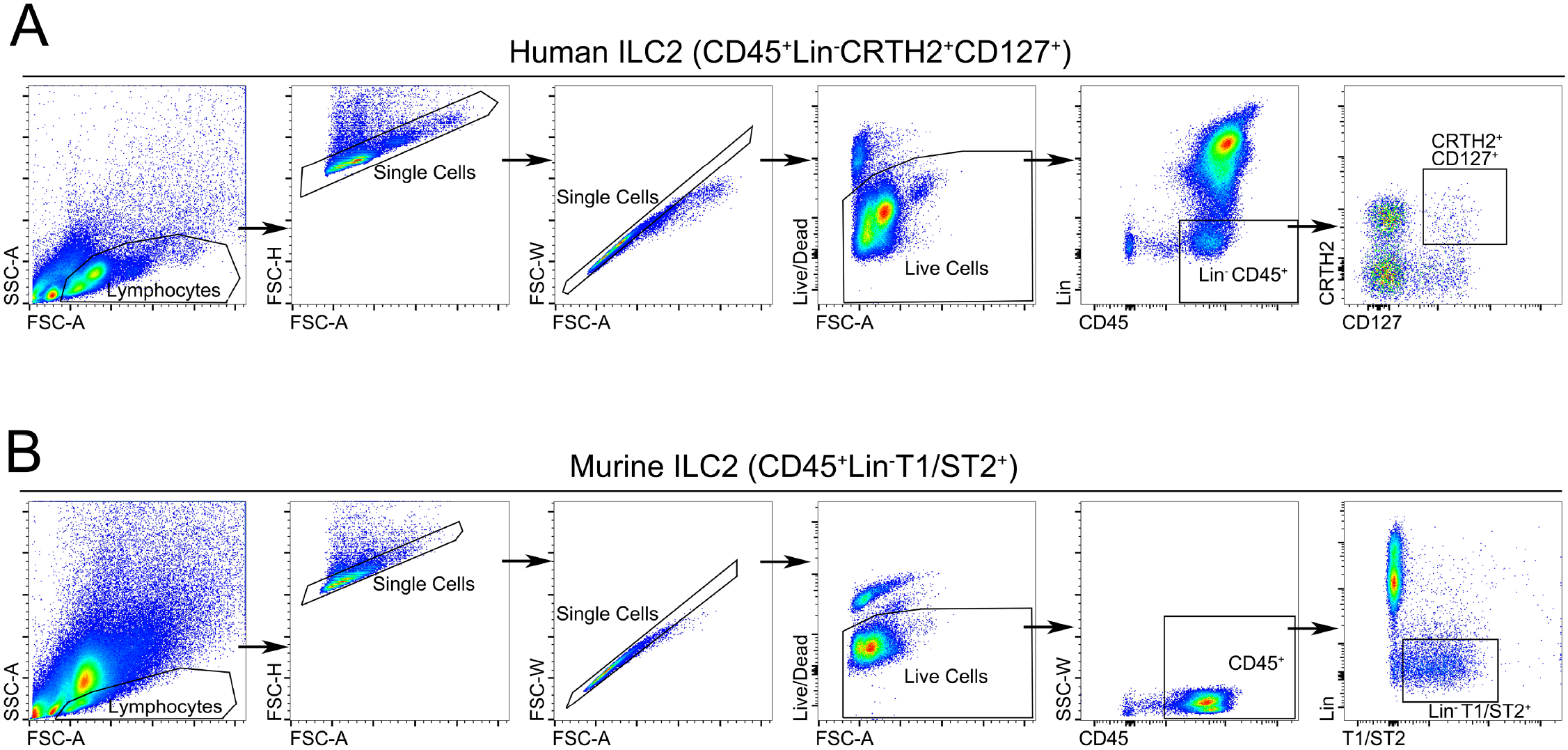
Human and murine ILC2 gating strategy. **A.** Human ILC2s were isolated from peripheral blood of healthy donors PBMCs or umbilical cord blood samples CBMCs and stained with antibodies against CD45 and lineage markers. Human ILC2s were sorted by the BD FACSAria cell sorter as CD45^+^Lin^-^CRTH2^+^CD127^+^ cells. **B.** Murine ILC2s were isolated from mouse lungs treated with recombinant IL-33 protein (250ng/mouse, *i.t.*) and stained with antibodies against CD45 and lineage markers as described in the Materials and Methods. Murine ILC2s were sorted by the BD FACSAria cell sorter as CD45^+^Lin^-^T1/ST2^+^ cells. The purity of sorted ILC2s was determined to be greater than 95%.

**Figure S2.**
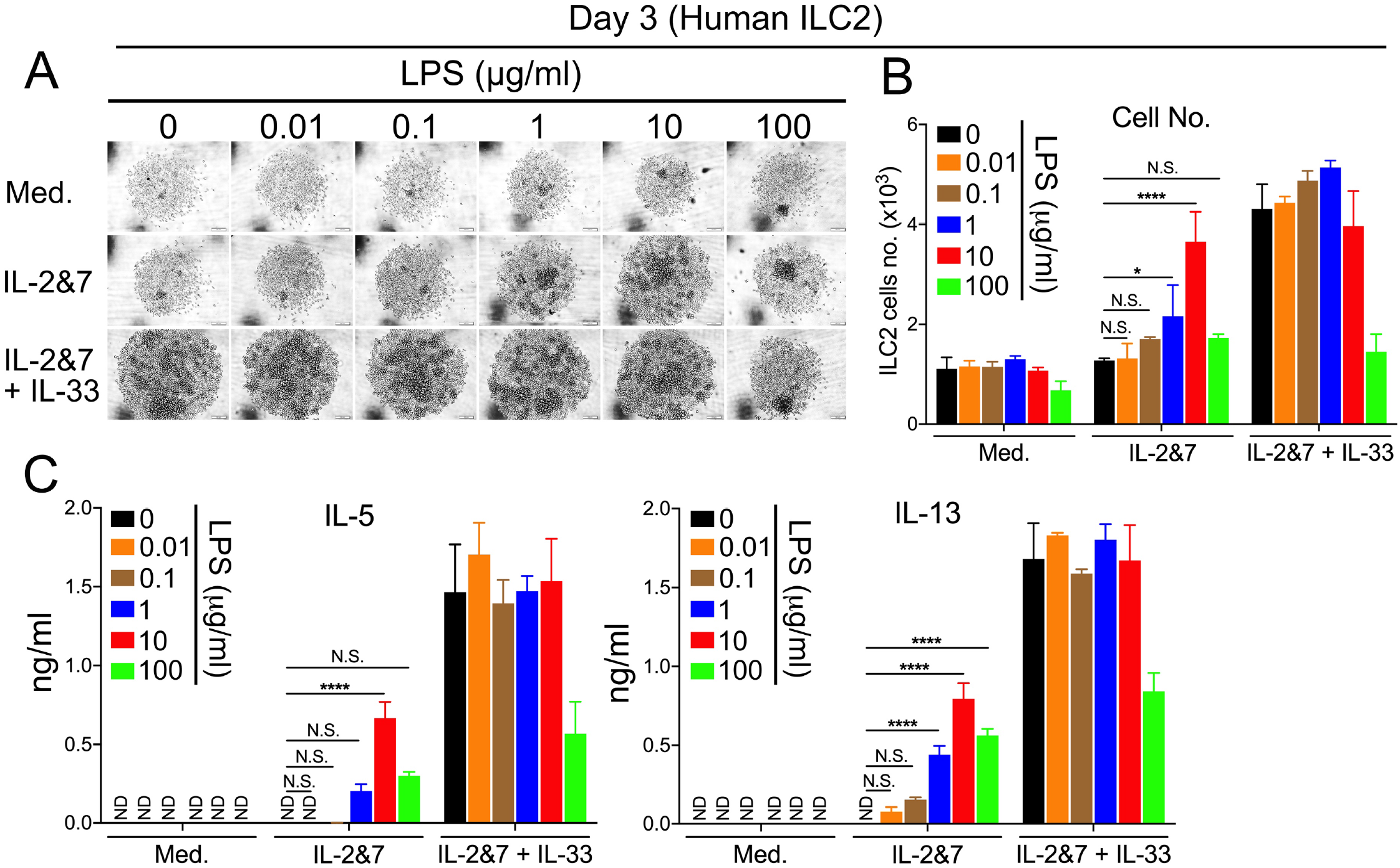
Analysis of LPS-stimulated human ILC2 cells on day 3. **A.** Light microscopic images showing the growth of ILC2 cells. FACS sorted human ILC2 cells were treated with the increased dose of LPS as indicated. Each image represented one well in which 1,000 cells were initially seeded. Scale bar, 100 μm. **B.** FACS showing the number of LPS-treated human ILC2 cells. **C.** ELISA measuring the production of IL-5 and IL-13 by human ILC2 cells treated with the increased dose of LPS. (P value≥0.05 was considered statistically insignificant, N.S., two-way ANOVA, * p < 0.05, **** p < 0.0001).

**Figure S3.**
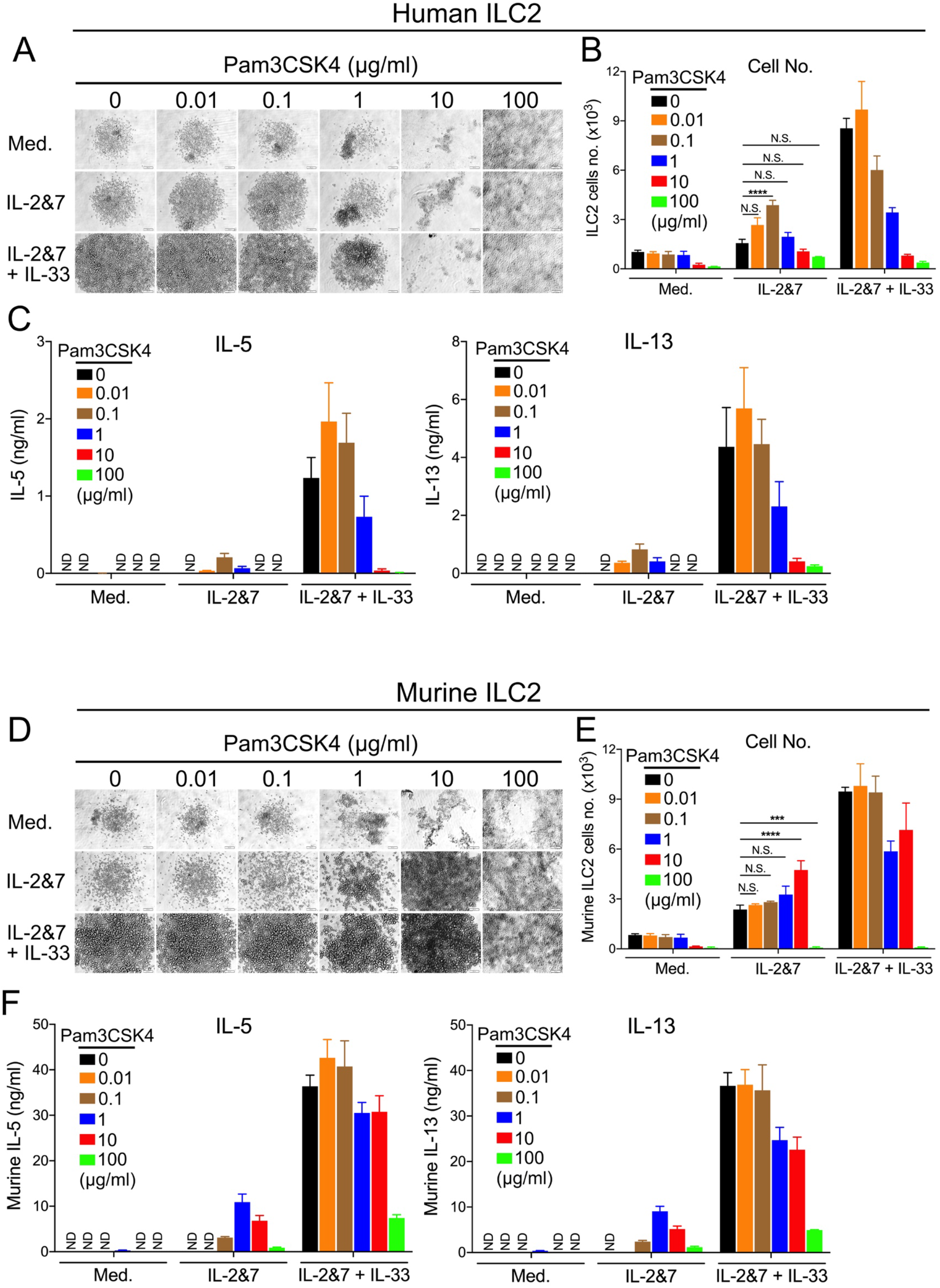
TLR2 agonist, Pam3CSK4, activates human or murine ILC2 cells at distinct concentrations. **A.** Light microscopic images showing the growth of human ILC2 cells stimulated with the increased doses of Pam3CSK4 as indicated. Each image represented one well in which 1,000 cells were initially seeded. Scale bar, 100μm. **B.** FACS showing the number of Pam3CSK4-treated human ILC2 cells. **C.** ELISA measuring the production of IL 5 and IL-13 by human ILC2 cells treated with the increased dose of Pam3CSK4. **D**. Light microscopic images showing the growth of murine ILC2 cells stimulated with the increased doses of Pam3CSK4 as indicated. **E.** FACS showing the number of Pam3CSK4 treated murine ILC2 cells. **F.** ELISA measuring the production of IL-5 and IL-13 by murine ILC2 cells treated with the increased dose of Pam3CSK4. (P value≥0.05 was considered statistically insignificant, N.S., two-way ANOVA, *** p < 0.001, **** p < 0.0001).

**Figure S4.**
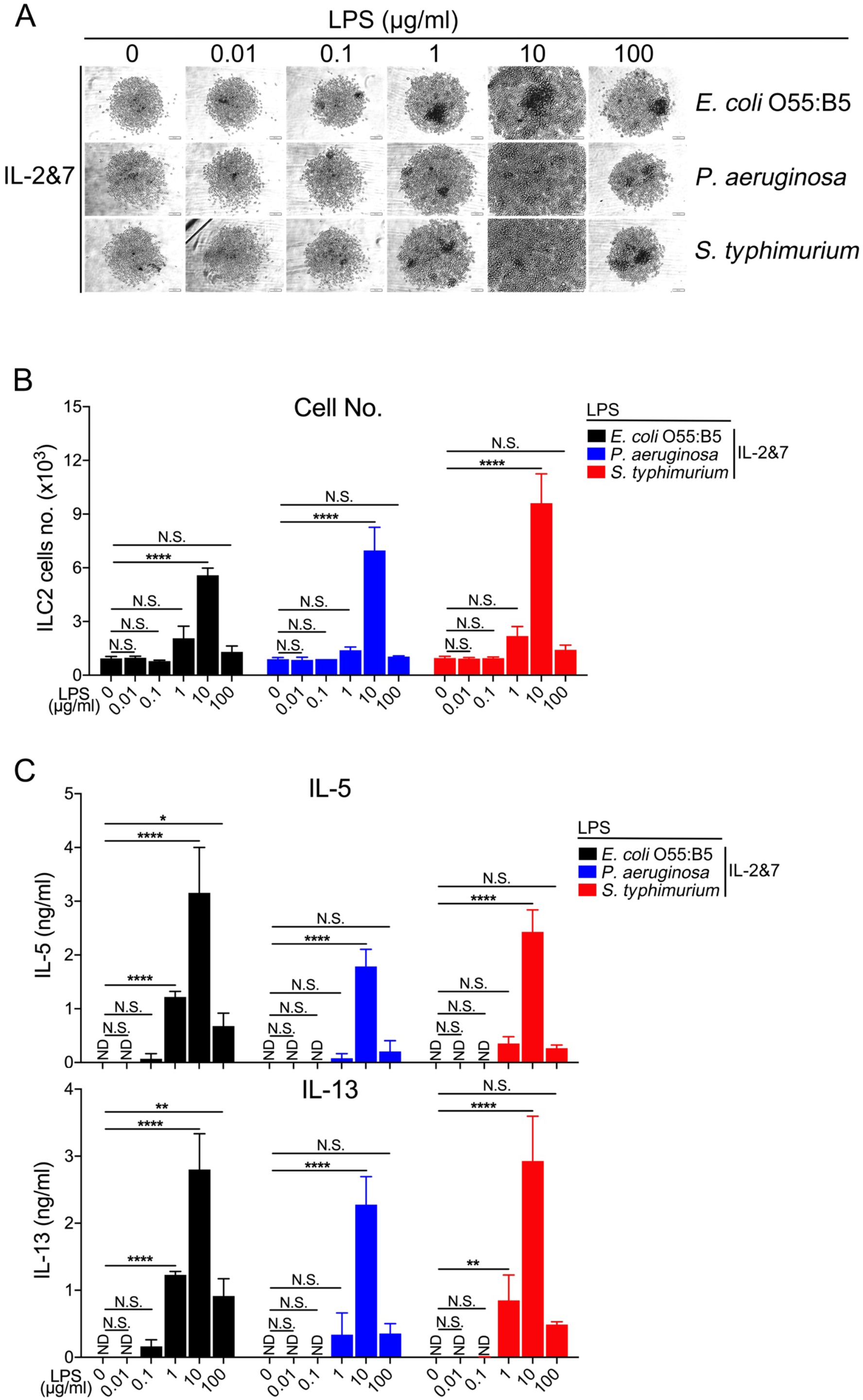
LPS from additional three Gram (-) bacterial species could stimulate the growth and cytokine production of human ILC2. **A.** Light microscopic images showing the growth of ILC2 cells treated with LPS isolated from *E. coli 055:B5*, *P. aeruginosa* and *S. typhimurium*. FACS sorted human ILC2 cells were treated with various LPS as indicated in a 96-well round-bottom plate for 5 days. Each image represented one well in which 1,000 cells were initially seeded. Scale bar, 100 μm. **B.** FACS showing the number of human ILC2 cells treated with various LPS as indicated. **C**. ELISA measuring the production of IL-5 and IL-13 by human ILC2 cells treated with various LPS as indicated. (P value≥0.05 was considered statistically insignificant, N.S., two-way ANOVA, * p < 0.05, ** p < 0.01, **** p < 0.0001).

**Figure S5.**
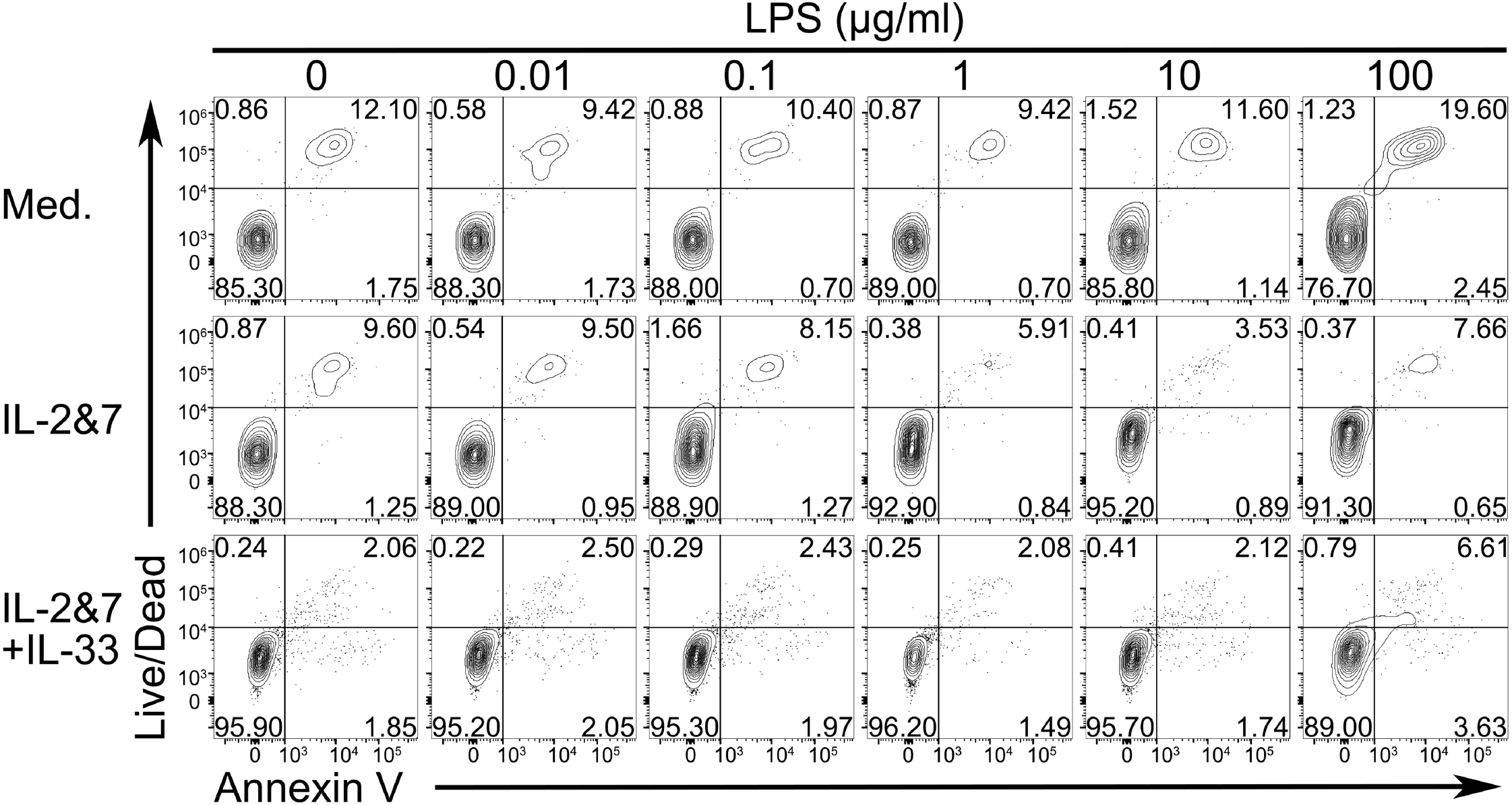
Cell death analysis of LPS-stimulated human ILC2 cells on day 3. FACS sorted human ILC2 cells were treated with the increased dose of LPS as indicated. FACS showing the cell death of LPS-treated human ILC2 cells.

**Figure S6.**
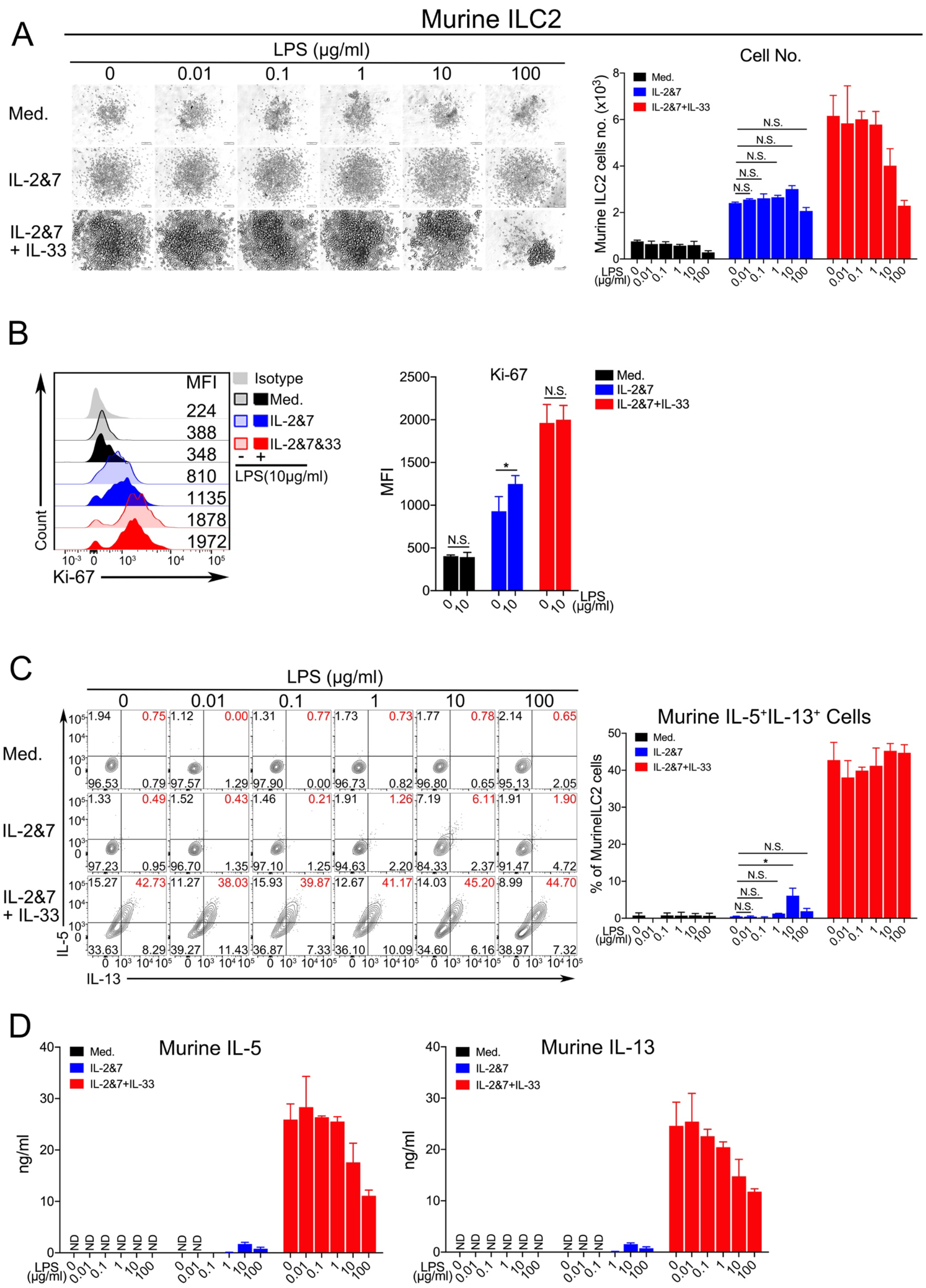
The effect of LPS on stimulating murine ILC2 cells is weak. **A.** Light microscopic images showing the growth of murine ILC2 cells. FACS sorted murine ILC2 cells were treated with the increased dose of LPS as indicated. Each image represented one well in which 1,000 cells were initially seeded. Scal bar, 100μm (left panel). FACS showing the number of LPS-treated murine ILC2 cells (right panel). **B.** The LPS-treated murine ILC2 cells were analyzed with Ki-67 staining. The median fluorescence intensity (MFI) and quantification of Ki-67 were shown. The result is a representative of two independent experiments. **C**. The representative gating strategy and percentage of IL5+IL13+-double positive murine ILC2 cells treated with the increased dose of LPS were analyzed by the intracellular staining. **D.** ELISA measuring the production of IL-5 and IL-13 by murine ILC2 cells treated with the increased dose of LPS. (P value≥0.05 was considered statistically insignificant, N.S., two-way ANOVA, * p < 0.05).

**Figure S7.**
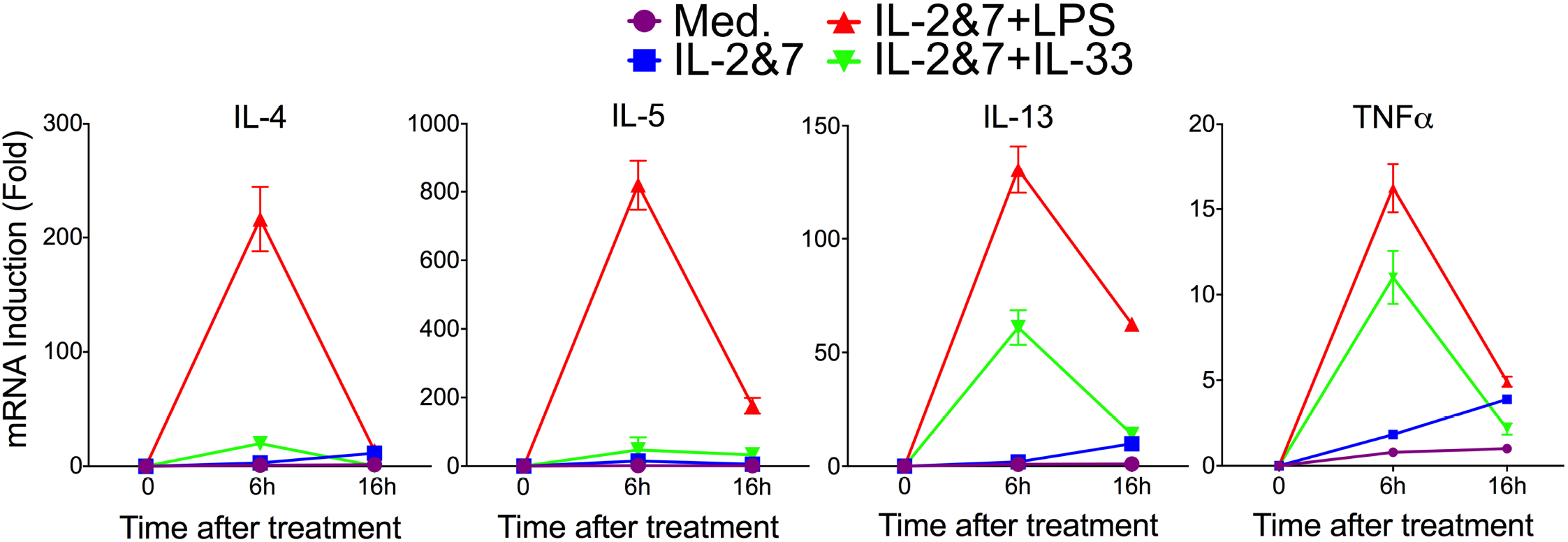
RT-qPCR analysis of selected gene expressions in LPS-stimulated human ILC2. Human ILC2 cells were collected at 0, 6 and 16 hours post LPS treatment. The mRNA expression of type 2 cytokines (IL-4, IL-5 and IL-13) and TNFα were measured with RT-qPCR.

**Figure S8.**
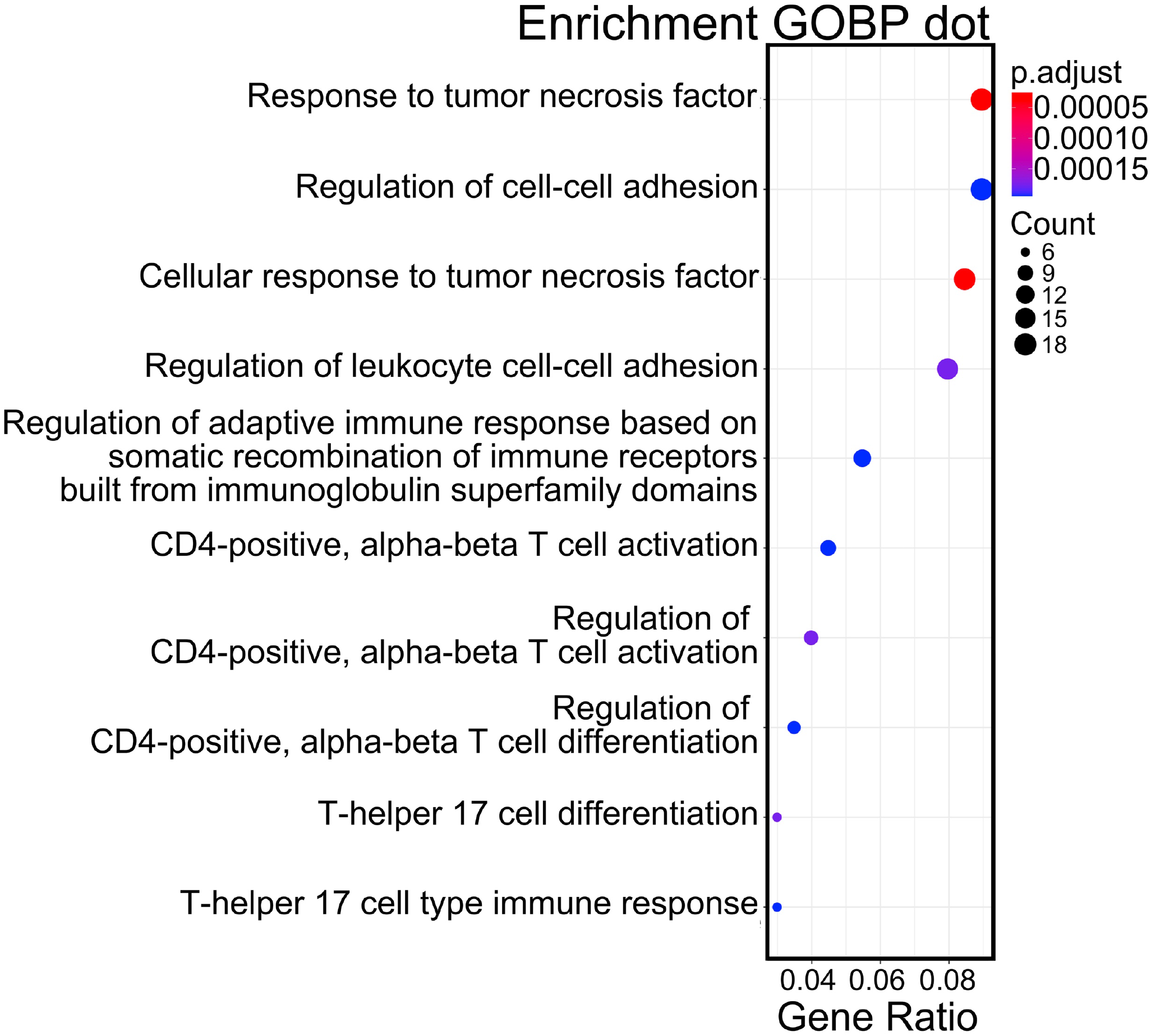
Dotplot of enrichment analysis with Gene Ontology biological process (GOBP).

